# Multi-organ functions of yolk sac during human early development

**DOI:** 10.1101/2022.08.03.502475

**Authors:** Rachel A Botting, Issac Goh, Antony Rose, Simone Webb, Justin Engelbert, Yorick Gitton, Emily Stephenson, Mariana Quiroga Londoño, Michael Mather, Nicole Mende, Ivan Imaz-Rosshandler, Dave Horsfall, Daniela Basurto-Lozada, Nana-Jane Chipampe, Victoria Rook, Pavel Mazin, MS Vijayabaskar, Rebecca Hannah, Laure Gambardella, Kile Green, Stephane Ballereau, Megumi Inoue, Liz Tuck, Valentina Lorenzi, Kwasi Kwakwa, Clara Alsinet, Bayanne Olabi, Mohi Miah, Chloe Admane, Dorin-Mirel Popescu, Meghan Acres, David Dixon, Rowen Coulthard, Steven Lisgo, Deborah J Henderson, Emma Dann, Chenqu Suo, Sarah J Kinston, Jong-eun Park, Krzysztof Polanski, Stijn Van Dongen, Kerstin B Meyer, Marella de Bruijn, James Palis, Sam Behjati, Elisa Laurenti, Nicola K Wilson, Roser Vento-Tormo, Alain Chédotal, Omer Bayraktar, Irene Roberts, Laura Jardine, Berthold Göttgens, Sarah A Teichmann, Muzlifah Haniffa

**Affiliations:** Biosciences Institute, Newcastle University, NE2 4HH, UK; Wellcome Sanger Institute, Wellcome Genome Campus, Hinxton, Cambridge CB10 1SA, UK; Sorbonne Université, INSERM, CNRS, Institut de la Vision, Paris, France; Department of Haematology, Wellcome-MRC Cambridge Stem Cell Institute, CB2 0AW, UK; MRC Laboratory of Molecular Biology, Cambridge Biomedical Campus, CD2 0QH, UK; Translational and Clinical Research Institute, Newcastle University, NE2 4HH, UK; NovoPath, Department of Pathology, Newcastle Hospitals NHS Foundation Trust, Newcastle upon Tyne, UK; Korea Advanced Institute of Science and Technology, Daejeon, South Korea; MRC Molecular Haematology Unit, MRC Weatherall Institute of Molecular Medicine, Radcliffe Department of Medicine, University of Oxford, OX3 9DS, UK; Department of Pediatrics, University of Rochester Medical Center, Rochester, 14642, NY, USA; Department of Paediatrics, University of Cambridge, Cambridge, UK; Department of Paediatrics, University of Oxford, OX3 9DS, UK

## Abstract

The yolk sac (YS) represents an evolutionarily-conserved extraembryonic structure that ensures timely delivery of nutritional support and oxygen to the developing embryo. However, the YS remains ill-defined in humans. We therefore assemble a complete single cell 3D map of human YS from 3-8 post conception weeks by integrating multiomic protein and gene expression data. We reveal the YS as a site of primitive and definitive haematopoiesis including a YS-specific accelerated route to macrophage production, a source of nutritional/metabolic support and a regulator of oxygen-carrying capacity. We reconstruct the emergence of primitive haematopoietic stem and progenitor cells from YS hemogenic endothelium and their decline upon stromal support modulation as intraembryonic organs specialise to assume these functions. The YS therefore functions as ‘three organs in one’ revealing a multifaceted relay of vital organismal functions as pregnancy proceeds.

**One Sentence Summary:** Human yolk sac is a key staging post in a relay of vital organismal functions during human pregnancy.

## Main Text

The primary human YS derives from the hypoblast at around the time of embryo implantation (Carnegie stage 4, CS4; ~1 post conception week (PCW)) (*1, 2*). A secondary YS beneath the embryonic disc supersedes the primary structure at around CS6 (~2.5PCW) and persists until 8PCW (*1, 2*). The secondary YS surrounds a vitelline fluid-filled cavity with three tissue compartments: mesothelium (a mesoderm-derived epithelial layer interfacing the amniotic fluid), mesoderm (which contains an array of cell types, including endothelial cells, blood cells and smooth muscle), and endoderm (interfacing the yolk sac cavity) (*1*). In phylogenetic terms, the YS is first seen in vertebrates with yolk-rich eggs e.g., birds, reptiles and amphibians, where its role is to extract macronutrients from yolk to sustain the embryo (*3*). The capacity to uptake, transport and metabolise nutrients is retained in mouse and human YS (*2*). Haematopoiesis originates in the YS in mammals, birds and some ray-finned fishes (*4*). The first wave of mouse YS haematopoiesis (primitive) yields primitive erythroid cells, macrophages and megakaryocytes from embryonic day 7.5 (E7.5) (*4, 5*). Following the onset of circulation, a second wave of erytho-myeloid and lympho-myeloid progenitors arise in the YS and supply the embryo (*6*). Finally, definitive haematopoietic stem cells arise in the aorta-gonad-mesonephros (AGM) region of the dorsal aorta and seed the fetal liver.

Limited evidence suggests that YS also provides the first blood cells during development in humans. Primitive erythroblasts expressing embryonic globin genes, surrounded by endothelium, emerge at CS6 (~2.5PCW) (*7, 8*). YS also produces megakaryocytes, mast cells and myeloid cells (*9*), although this has not yet been evidenced directly in functional studies. Transplantation of human developmental tissues into immunodeficient mice has pinpointed the origin of definitive haematopoietic stem and progenitor cells (HSPCs) (defined as long-term multilineage repopulating cells) within the AGM region of embryo at CS14 (~5PCW) (*10*). Equivalent cells are subsequently found in the YS at CS16 and in the liver from CS17 (*10*). This sequence was also documented by following the transcriptional signature of definitive HSPCs across organs (*11*). While the process of definitive HSPC emergence from hemogenic endothelium (HE) has been reconstructed in human AGM (*11, 12*), the process by which earlier progenitors arise in human YS has not been studied. Several key questions about human YS haematopoiesis remain unanswered: what is the full repertoire of human YS-derived blood cells, does the YS produce limited progenitors or HSPCs, do YS progenitors/HSPCs contribute to long-lived populations such as tissue-resident macrophages, do YS progenitors/HSPCs arise from HE, and what are the extrinsic regulators of this process.

In this study we report a time-resolved atlas of the human YS combining single cell multiomics with 3D light-sheet microscopy and multiplex RNA *in situ* hybridisation, providing the first comprehensive depiction of the metabolic and haematopoietic functions of the human YS, as well as a benchmark for *in vitro* culture systems aiming to recapitulate early human development.

## Results

### A single cell atlas of the human yolk sac

We performed droplet-based scRNA-seq profiling of human YS, including both membrane and contents, and integrated with external datasets to yield 169,798 high quality cells from 10 samples spanning 4-8PCW (CS10-CS23), which can be interrogated on our interactive web portal (https://developmental.cellatlas.io/yolk-sac; password: ys2022) (*13*) (**Fig. 1A-C, S1A-C, Table S1-5**). All datasets used for cell state validation, *in vitro* iPSC culture and crossspecies comparisons are shown in **Fig. 1A (Table S6-7**).

**Fig. 1:**
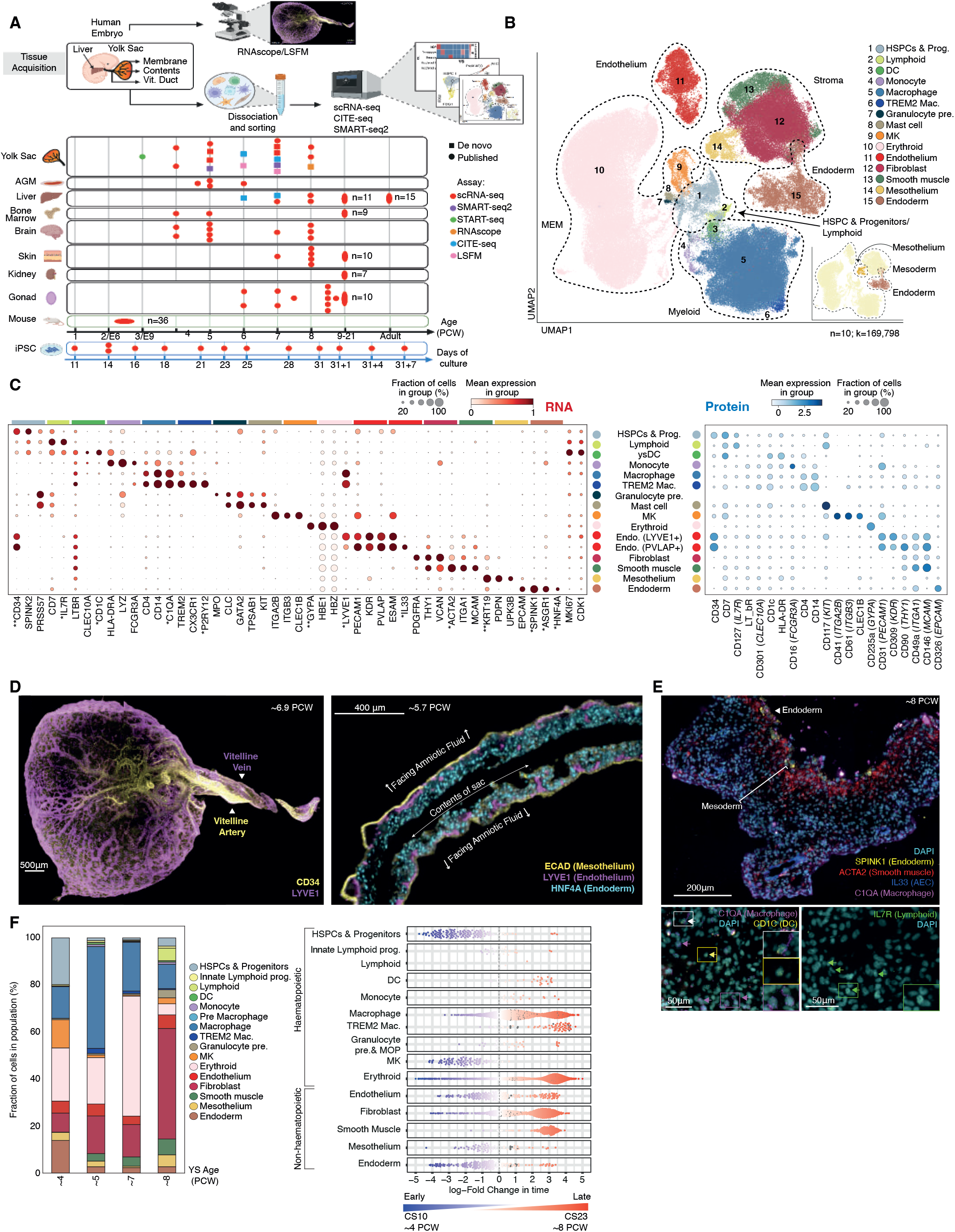
A single cell atlas of the human yolk sac. **(A)** Summary of data included in analyses. Squares represent data generated for this study and circles represent published datasets: YS (*9, 11, 45, 53*), AGM (*11*), liver (*9*), fetal BM (*54*), fetal brain (*44*), fetal skin (*45*), fetal kidney (*55*), fetal gonads (*46*), mouse (*56*), iPSC (*11, 21*). Shape colour indicates the assay used to generate data (**Table S6**). (**B**) UMAP visualisation of YS scRNA-seq data (n=10 independent biological replicates; k=169,798), with colours representing broad cell states: DC= dendritic cell, Mac= macrophage, MEM= megakaryocyte-erythroid-mast cell lineage, MK= megakaryocyte, pre.= precursor. Insert shows the same UMAP coloured by YS tissue layer (**Table S5**). (**C**) Left: Dot plot showing the expression level (by colour) and percent expression (by dot size) of broad cell state-defining genes in scRNA-seq data as shown in **b,** with data scaled to a maximum value=1. Right: Dot plot showing equivalent protein expression (by colour) and percent expression (by dot size) of broad cell states from n=2 biologically independent YS CITE-seq samples, with data scaled zero_centre=False. * indicates genes validated by RNAscope and ** indicates equivalent proteins validated by IHC/IF (**Table S4**). (**D**) Light-sheet fluorescence microscopy images of YS. Left: CD34^+^ and LYVE1^+^ vascular structures (representative ~6.9PCW sample; scale bar=500μm; **Movie S1-2**). Right: Location of LYVE1^+^ vascular structures within HNF4A^+^ endoderm and adjacent to ECAD^+^ mesothelium (representative ~5.7PCW sample; scale bar=400μm). (**E**) RNAscope images of YS, showing a representative 8PCW sample. Top: demonstrating endoderm (*SPINK1*), smooth muscle (*ACTA2*), AEC (*IL33*) and macrophages (*C1QA*) (scale bar=200μm). Bottom: demonstrating DCs (*CD1C;* yellow arrow), macrophages (*C1QA;* magenta arrow), mac-DCs (*CD1C^+^C1QA^+^;* white arrow) and lymphoid cells (*IL7R;* green arrow) (scale bar=50μm). (**F**) Left: Bar graph showing the proportional representation of broad cell states to YS scRNA-seq data grouped by gestational age in PCW. Right: Milo beeswarm plot showing differential abundance of YS scRNA-seq neighbourhoods across time, where blue neighbourhoods are significantly enriched (SpatialFDR<0.1,logFC<0) early in gestation (CS10-11), red neighbourhoods are enriched later (CS22-23) (SpatialFDR<0.1,logFC>0) and colour intensity denotes degree of significance. Abbreviations: as per **(b)** and MOP= monocyte progenitor and pre= precursor (**Table S19**).

With iterative graph-based leiden clustering and annotation, the integrated YS scRNA-seq dataset yielded 43 cell types, which we grouped into 15 broad categories including haematopoietic cells, endoderm, mesoderm and mesothelium (**Fig. 1B-C, S1B-C, Table S3-5**). We consistently apply the term HSPC for cells expressing a core HSPC signature e.g. *CD34, SPINK2, HLF,* without implying long-term repopulating capacity or multilineage potential. With comparison datasets, unless otherwise specified, we adopt published annotations (**Table S6-7**). We demonstrate the key marker genes for these broad cell categories, validated by plate-based scRNA-seq (**Fig. S1D-E, Table S8, S5**) and surface protein expression from CITE-seq analysis of n=2 YS cell suspension (**Fig. 1C, S1F-G, Table 9)**. We used surface protein expression data to construct a decision tree that identified combinatorial antibodies deployable for YS cell type purification and functional characterisation (**Fig. S2A**). FACS isolation of CD45^-^ cells with high scatter and Smart-seq2 analysis enabled enrichment and validation of YS endoderm, mesothelium and erythroid cells (**Fig. S1D-E, Table S8, S5**). We generated matched embryonic liver scRNA-seq data for established and late stage YS at CS18 and CS22-23 respectively (n=3; n=2 previously reported (*9*)) (**Fig. S2B-F**), validated by liver CITE-seq data (**Fig. S3A-C, Table S10**), which confirmed the presence of pre-macrophages only in YS and discrete B-cell progenitor stages only in the liver (**Fig. S1C, S3C)**. Around half of YS lymphoid cells were progenitors, which terminated in NK and ILC precursor states on force directed graph (FDG) visualisation (**Fig. S3D**). A small population of cells were termed ‘B lineage’ due to expression of *CD19, CD79B* and *IGLL1.* They did not express genes typical of B1 cells (*CCR10, CD27, CD5*). Given the absence of distinct B cell progenitor stages and their later emergence (>5PCW), these may constitute migratory B cells of fetal liver origin (**Fig. S3D-E**).

To facilitate future use of our YS atlas, we employed a low-dimensional logistic regression (LR) framework (see Methods). The trained LR models and weights from trained scVI models are provided via our interactive web portal to enable mapping of scRNA-seq datasets using CellTypist (*14*) and transfer learning with single-cell architectural surgery (scArches) (*15*), respectively. Corresponding cross-tissue projection probability matrices are provided as Supplementary Tables (**Table S11-18**).

Using selected marker genes and proteins from our droplet-based datasets (**Fig. 1C**), we performed 3D visualisation of YS by light sheet microscopy, demonstrating the CD34^hi^PVLAP^hi^LYVE1^l^° vitelline artery and a CD34^lo^PVLAP^lo^LYVE1^hi^ vitelline vein contiguous with a branching network of CD34^-^PVLAP^lo^LYVE1^hi^ vessels within the YS (**Fig. 1D, Fig. S3F**). The LYVE1^hi^ vessels localised within HNF4A^+^ endoderm and adjacent to the ECAD^+^ mesothelium (**Fig. 1D, S3G-I**). Rare *IL7R^+^* lymphoid cells, and *CD1C^+^C1QA^+/^* dendritic cells (DCs) were identified within the mesoderm, while *ACTA2*^+^ smooth muscle cells surrounded *IL33^+^* vessels, forming a sub-layer between mesoderm and *SPINK1^+^* endoderm (**Fig. 1E, S4A-B**).

To investigate the functional relevance of changes in YS cell composition during development, we next assessed proportional representation of cell states by gestation. In early YS (CS10; ~4PCW) HSPCs, erythroid cells, macrophages and megakaryocytes (MK) were the most prevalent cell types with both HSPCs and MKs proportionately diminished thereafter, while production of erythroid cells and macrophages was sustained. DC and *TREM2^+^* microglia-like cells did not emerge until >6PCW (**Fig. 1F, Table S4**). The ratio of haematopoietic to non-haematopoietic cells was around 3:1 in early YS (CS10; ~4PCW), but with expansion of fibroblasts particularly, the ratio in late YS (CS22-23; ~8PCW) approached 1:3 (**Fig. 1F**). We performed graph-based differential abundance testing with Milo (*16*) to assess heterogeneity within cell states by gestation. Both erythroid cells and macrophages had early and late-gestation specific molecular states but MKs were transcriptionally homogenous across gestation (**Fig. 1F, Table S19**). Early gestation molecular states were characterised by significantly upregulated ribosomal and glycolytic genes (e.g., *RPS29, ENO1, LDHB*), suggesting a common early burst of translation (**Fig. 1F, S4C, Table S4**). In contrast, cells at later gestation upregulated specific lineage-defining genes compared to their earlier counterparts (**Fig. 1F**).

### Multi-organ functions of YS

To probe the nutritional/metabolic functions of YS, we next focused our scRNA-seq analysis on the non-haematopoietic cells. We identified an endoderm cell state co-expressing *APOA1/2, APOC3* and *TTR,* similar to embryonic/fetal hepatocytes, which was present from gastrulation (**Fig. 2A, S4D, Table S3, S20, S7**). Compared with embryonic liver hepatocytes, YS endoderm expressed higher levels of transcripts for serine protease 3 (*PRSS3*), Glutathione S-Transferase Alpha 2 (*GSTA2*) and multi-functional protein Galectin 3 (*LGALS3*), while embryonic liver hepatocytes expressed a more extensive repertoire of detoxification enzymes, including alcohol and aldehyde dehydrogenases (*ADH1A, ALDH1A1*) and cytochrome P450 enzymes involved in metabolism of steroid hormones and vitamins (*CYP3A7*), fatty acids (*CYP4A11*) and the conversion of cholesterol to bile acids (*CYP27A1*) (**Fig. S4D, Table S20**). YS endoderm and embryonic liver hepatocytes shared gene modules implicated in coagulation and lipid and glucose metabolism (**Fig. 2B, Table S21**). These modules were also conserved between human and mouse extraembryonic endoderm (**Fig. S4E-F, Table S21**). We validated *in situ* the expression of lipid transport (alpha-fetoprotein, albumin and alpha-1-antitrypsin) and coagulation proteins (fibrin) in YS endoderm and embryonic liver hepatocytes (**Fig. 2C, D**).

**Fig. 2:**
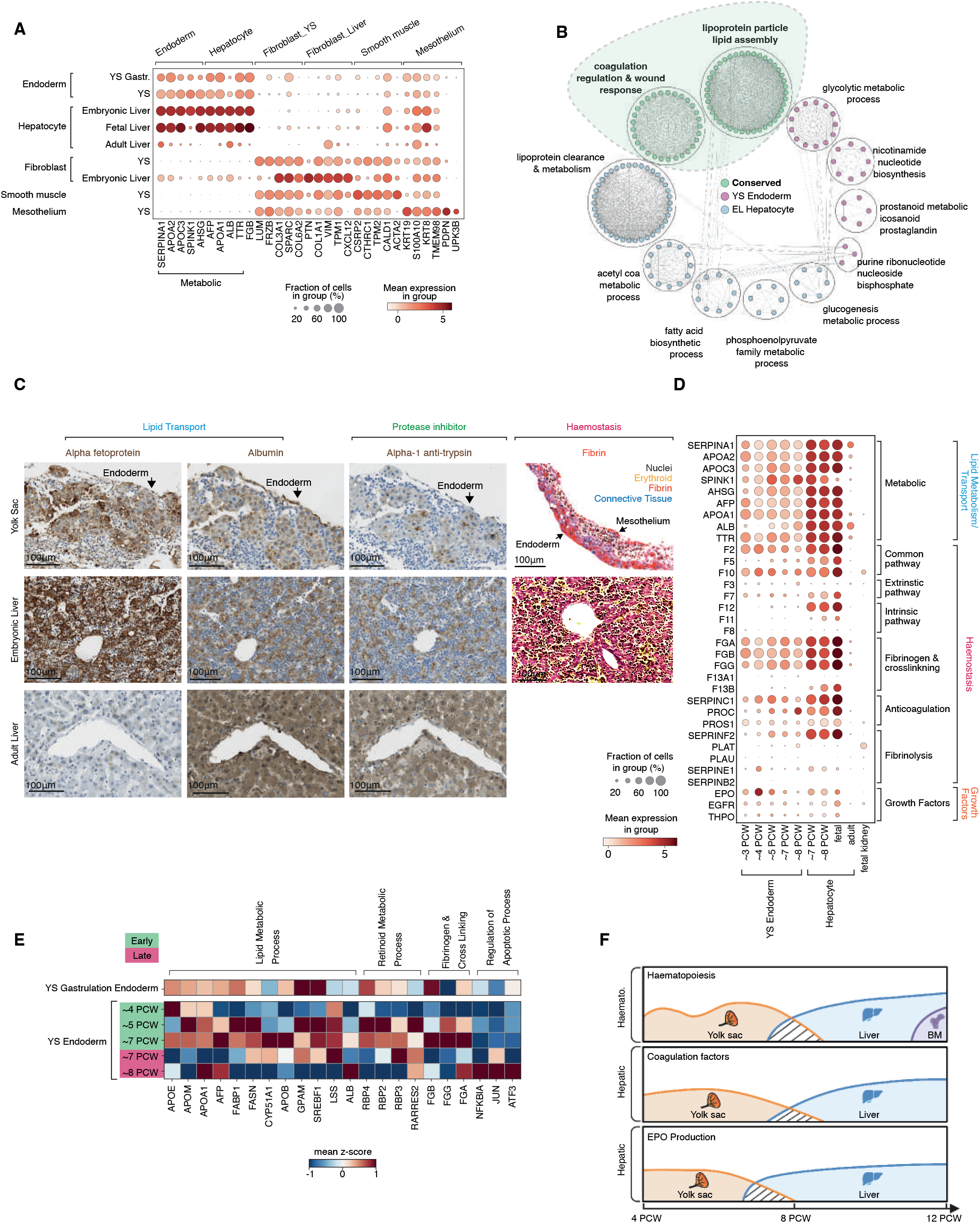
Multi-organ functions of YS. **(A)** Dot plot showing the expression level (colour scale) and percent expression (dot size) of DEGs in YS (main and gastrulation (gastr.) data) stromal cell states, matched embryonic, fetal and adult liver stromal cell states and iPSC stromal cell states (*21*) (**Table S3, 20, 7**). All datasets independently scaled to max value=10 and then combined, except YS and matched EL scRNA-seq data which were scaled together. Genes involved in endoderm metabolic function are grouped. **(B)** Flower plot of the significant pathways upregulated in YS endoderm (pink), embryonic liver (EL) hepatocytes (blue) and conserved across both tissues (green). Lines indicate connected nodes of expression (**Table S21**). **(C)** Left and middle: IHC staining of alpha fetoprotein (*AFP*), albumin (*ALB*) and alpha-1 antitrypsin (*SERPINA1*) in 8PCW YS, 8PCW EL and healthy adult liver. Representative images from 1 of n=5 biological independent YS (4-8PCW), 1 of n=3 biologically independent ELs (7-8PCW) and 1 of n=3 biologically independent adult livers. Scale bar=100μm. Right: MSB-stained 8PCW EL (representative of n=3 biologically independent samples) and 4PCW YS (representative of n=3 biologically independent samples). Nuclei (grey), erythroid (yellow), fibrin (red), and connective tissue (blue). See ‘Immunohistochemistry’ section in Methods for details regarding pseudo-colouring shown. Scale bars=100um. **(D)** Dot plot showing the expression level (colour scale) and percent expression (dot size) of haemostasis factors expressed by YS endoderm (main and gastrula data), embryonic, fetal and adult liver hepatocytes, and endoderm from fetal kidney (*55*). Grouped by pathway and role. All datasets independently scaled to max value=10 and then combined, except YS and matched EL scRNA-seq data which were scaled together. **(E)** Matrix heatmap of Milo-generated DEGs in endoderm from YS scRNA-seq including gastrulation datasets (**Table S19**). DEGs are grouped by function. Each dataset scaled zero centre=False and YS endoderm standard_scale=‘var’. **(F)** Schematic of the relative contribution of YS (orange), EL (blue), BM (purple) to haematopoiesis, coagulation factors and EPO synthesis in the first trimester of human development.

We further explored the gene module ‘coagulation regulation’, conserved between YS and liver, (**Table S21**) by examining expression of pro- and anticoagulant proteins across samples, grouped by gestational age (**Fig. 2D**). From the earliest timepoints, YS endoderm expressed components of the *F3* (Tissue factor)-activated coagulation pathway, particularly *F2* (Thrombin) and *F10* (Factor X) and anticoagulant proteins *SERPINC1* (Antithrombin III) and *PROS1* (Protein S) (**Fig. 2D**). Tissue factor, Antithrombin III and Fibrinogen subunits were also expressed in mouse extraembryonic endoderm (**Fig. S4E**). Factors for the intrinsic pathway (triggered by exposed collagen) were minimally expressed in YS, but were expressed by embryonic liver hepatocytes (**Fig. 2D**). Given the importance of Tissue factor in YS angiogenesis (*17, 18*), it is likely the coagulant and anticoagulant pathways develop in parallel as a means to balance haemostasis.

In addition to their metabolic and coagulation functions, YS endoderm cells expressed EPO and THPO that are critical for erythropoiesis and megakaryopoiesis (**Fig 2D, S4G**). EPO is known to be produced by mouse YS endoderm (*19*), and both growth factors are produced by human FL hepatocytes (*9*), but their combined presence has not previously been reported in human YS.

We next investigated how multi-organ functions of YS endoderm change over time. Milo-generated DEGs between early and late YS endoderm neighbourhoods revealed active retinoic acid and lipid metabolic processes until 7PCW, after which genes associated with cell stress and death were expressed (**Fig. 2E, Table S22**). Collectively, our findings describe a critical role of the human YS to support haematopoiesis, metabolism, coagulation and erythroid cell mass regulation, prior to these functions being taken over by fetal liver, and ultimately, by adult liver (metabolism and coagulation), bone marrow (haematopoiesis) and kidney (erythroid cell mass regulation) (**Fig. 2F**).

### Primitive versus definitive haematopoiesis in YS and liver

Whether the YS is a site of definitive haematopoiesis in humans has not been fully resolved (*20*). Human YS progenitors spanned two groups-HSPC characterised by *SPINK2, CYTL1* and *HOXB9,* and cycling HSPC characterised by cell-cycle associated genes such as *MKI67* and *TOP2A* (**Fig. S5A**). HSPC and cycling HSPC had the highest probability of class prediction to fetal/embryonic liver HSC and MPP respectively (**Fig. S5B**). We further identified HSPC sub-states expressing markers characteristic of primitive (*DDIT4, SLC2A3, RGS16, LIN28A*) and definitive (*KIT, ITGA4, CD74, PROCR*) HSPCs in human YS/AGM (*11*), present in both HSPC and cycling HSPC fractions (**Fig. 3A-C**). Both the primitive and definitive HSPC sub-states expressed canonical HSPC genes such as *SPINK2, HOPX* and *HLF* but diverged in expression of genes involved in multiple processes such as enzymes (*GAD1*), growth factors (*FGF23*), adhesion molecules (*SELL*) and patterning genes (*HOXA7*) (**Fig. 3A, S5C**). Differential protein expression in CITE-seq data indicated that CD194 (CCR4), CD357 and CD122 mark primitive HSPCs while CD197 (CCR7), CD193 and CD48 are preferentially expressed on definitive HSPCs (**Fig. S5D**). We confirmed that an iPSC-derived culture system reported to generate definitive HSPCs did express markers characteristic of definitive HSPCs (*11*), but an iPSC-derived culture system optimised for macrophage production (*21*) did not (**Fig. 3A**). To assess cross tissue HSPC heterogeneity, we integrated HSPCs across hematopoietic organs (see Methods) (**Fig. 3C, S5E)**. By kernel density estimation in integrated UMAP embeddings, YS definitive HPSCs co-localized with definitive HSPCs from age-matched liver (**Fig. 3C, S5E).** From exclusively primitive HSPCs at ~3PCW, we observed rapid accumulation of definitive HSPCs after AGM development CS14 (~5PCW), likely accounting for the increase in YS HSPC/progenitor fraction at 8PCW (**Fig. 1F, 3B, S5F**).

**Fig. 3:**
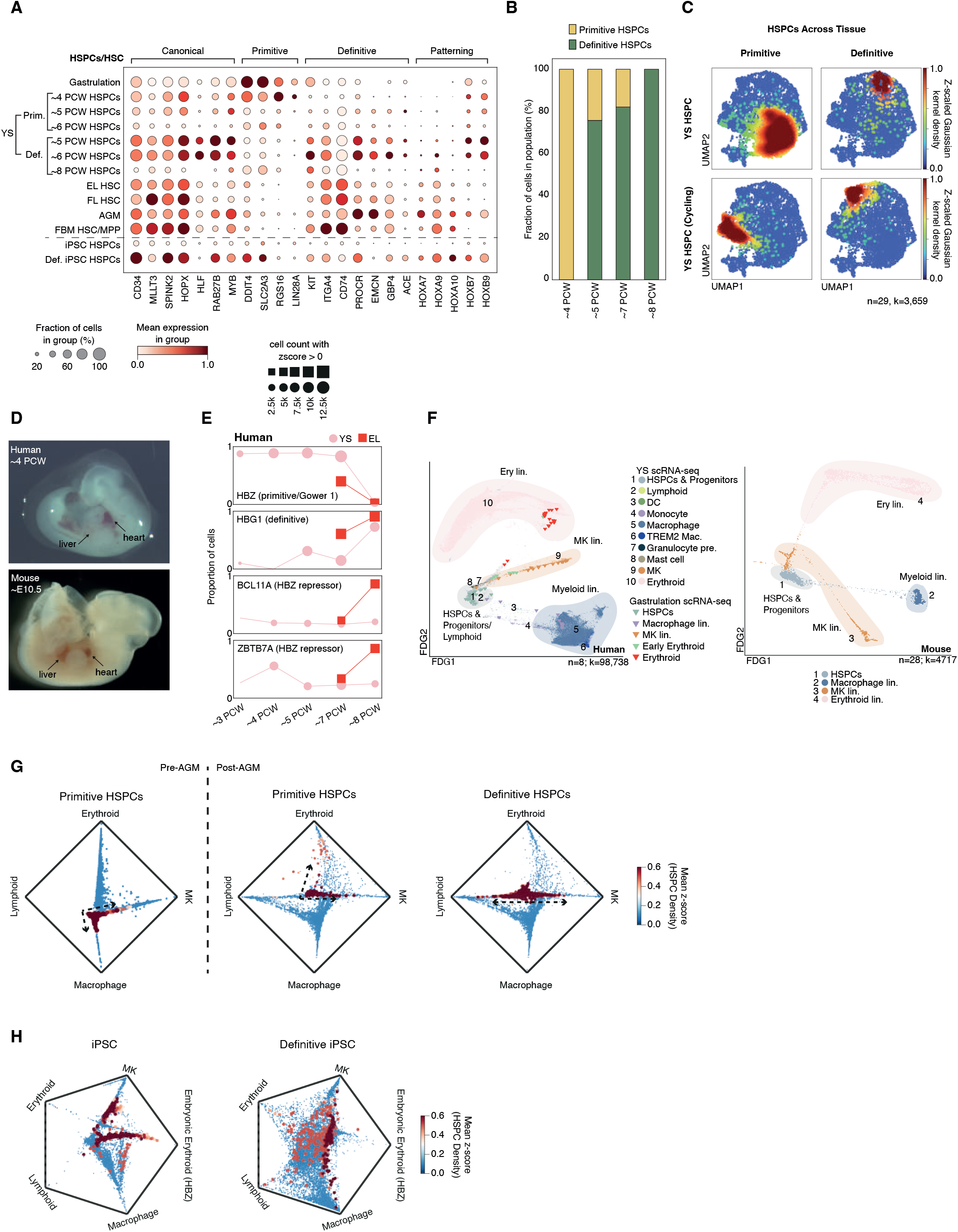
Primitive versus definitive haematopoiesis in YS and liver. **(A)** Dot plot showing the mean variance scaled expression level (colour scale) and percent expression (dot size) of canonical, primitive and definitive, and patterning HSC markers expressed between YS HSPCs, including gastrulation (*57*), AGM HSPC (*58*), matched EL HSC, FL HSC (*9*), fetal BM HSC/MPP (*54*), iPSC-derived HSPC (*21*) and definitive iPSC-derived HSPC (*11*). **(B)** Bar graph showing the proportional representation of primitive YS HSPCs to definitive YS HSPCs in the main YS scRNA-seq data (grouped by gestational age in PCW). **(C)** Density plots showing the distribution of YS HSPC (top), cycling HSPC (bottom) with primitive (left) and definitive signatures (right) in the integrated UMAP landscape of HSPC/HSCs from YS including gastrulation (n=10, k=2,597), AGM (*58*) (n=1, k=28), matched embryonic liver (EL) (n=3, k=412), fetal liver (FL) (*9*), fetal bone marrow (*54*) (FBM) (n=9, k=92), iPSC-derived HSPC (*21*) and definitive iPSC-derived HSPC (*11*) scRNA-seq datasets. Colour of HSC/HSPC cells represents the z-scored kernel density estimation (KDE) score for each population (**Table S5**). **(D)** Image of a ~4PCW/CS12 human embryo (top; representative from n=4 biologically independent samples) and ~E10.5/CS12 mouse embryo (bottom; n=1) with the heart and liver labelled. **(E)** Line graphs showing the relative change in proportion of erythroid lineage cells (y-axis) enriched in expression of globins over gestational age, including *HBZ, HBG1* and HBZ-repressors *BCL11A* and *ZBTB7A.* Pink lines= human YS main and gastrulation scRNA-seq data. Red lines= matched scRNA-seq data. Globins are grouped by their role in primitive haematopoiesis, definitive haematopoiesis and repression. **(F)** Force directed graph (FDG) visualisation of haematopoietic cell states in the YS scRNA-seq dataset (n=8, k=98,738; dots) integrated with human gastrulation (*57*) scRNA-seq dataset (n=1, k=91; triangles) (left), and equivalent cell states found in the mouse gastrulation scRNA-seq dataset (*56*) (n=28, k=4,717; dots) (right). Colours represent cell states and clouds mark lineages **(G)** Circular plots showing relative absorption probabilities of lineage-state transition between primitive HSPCs in the YS pre-AGM (CS10-11) (left) and primitive and definitive HSPCs in the YS post-AGM (>CS14) (right). Colour indicates the HSPC population density as a z-scored kernel density estimation (KDE) score and the position of HSPC population densities indicate respective lineage priming probability between Macrophage, lymphoid (NK and B lineage), erythroid and MK terminal states. **(H)** Circular plots showing relative absorption probabilities of lineage-state transition between iPSC-derived HSPCs (left) and definitive iPSC-derived HSPCs (right). Colour indicates the HSPC population density as a z-scored kernel density estimation (KDE) score and the position of HSPC population densities indicate respective lineage priming probability between Macrophage, lymphoid (NK and B lineage), erythroid (any erythroid cell with individual Z-score of HBA1,HBA2,HBG1,HBG2,HBD > 0), embryonic erythroid (any erythroid cell with HBZ Z-score > 0) and MK terminal states.

Next, we aimed to establish when the liver takes over from YS haematopoiesis and whether it initially uses primitive or definitive HSPCs. Prior to AGM (and to circulation), the human embryonic liver is macroscopically pale, suggesting that erythropoiesis predominantly occurs in the YS (**Fig. 3D**). We tracked the proportional representation of haemoglobin (Hb) subtypes over time as a proxy for YS versus embryonic liver contributions, as the zeta globin chain (*HBZ*) is restricted to primitive erythroblasts while definitive erythroblasts in fetal liver express gamma globins (*HBG1*) (*22, 23*) (**Fig. 3E)**. Our observation of sustained *HBZ* production in YS, for several days prior to liver bud formation (4 PCW), is consistent with a scenario where YS supports sustained erythropoiesis. The minimal zeta globin expression and expression of *HBZ* repressors by embryonic liver erythroblasts suggests that embryonic liver erythropoiesis is predominantly definitive, as we have previously shown (*9*), and that the transition to liver-dominated erythropoiesis occurs around 8PCW (**Fig. 3E**). This differs from the situation in mice, whereby immature definitive erythroid progenitors exit the YS and rapidly mature in other sites (*24*), as evidenced by the macroscopically red appearance of the mouse embryonic liver prior to AGM maturation (**Fig. 3D**). Tracking Hb subtype usage in the mouse, we noted two waves of erythropoiesis pre-AGM: primitive haematopoiesis in the YS (initially *Hbb BH1* and *Hba X-Hba1*) and pro-definitive haematopoiesis mirrored in both YS and torso (*Hbb BT1* and *Hbb BS*) (**Fig. S5G-I**).

Given the co-existence of primitive and definitive HSPCs in most of our YS samples, we examined an earlier reference (prior to AGM-HSPC derivation) to explore the differentiation potential of primitive HSPCs. At gastrulation (~3PCW), the YS haematopoietic landscape had a tripartite differentiation structure, with erythroid, MK and myeloid differentiation (**Fig. 3F)**. This structure was also observed in mouse YS **(Fig. S6A-B, Table S13, S5**). A differential-fate-prediction analysis demonstrated that primitive HSPCs pre-AGM at CS10-11 (~4PCW) were biased towards myeloid cell fates, however the abundance of differentiating erythroid and MK cells at this point suggests that an earlier wave of erythroid/MK production occurred (**Fig. 3G, S6C**). Post-AGM, the model predicted that remaining primitive HSPCs were erythroid/MK-biased while definitive HSPCs were biased towards lymphoid and MK fates (**Fig. 3G**). This is in keeping with the first appearance of YS lymphoid cells (ILC progenitors and NK cells, and B lineage cells) post CS14 (**Fig. 1F**). Differential-fate-prediction analyses revealed that iPSC-derived HSPCs were embryonic erythroid (i.e. erythroid cells expressing *HBZ*), myeloid and MK-biased whilst definitive iPSC-derived HSPCs were lymphoid, MK, non-embryonic erythroid and myeloid-primed, consistent with the predicted lineage potential of their *in vivo* primitive and definitive counterparts (**Fig. 3H, S6D**).

### The lifespan of YS HSPCs

Following predictions that primitive HSPC differentiation potential changes over time, we sought to examine the extrinsic support received by HSPCs across their lifespan. HSPCs arise from hemogenic endothelium (HE) in the aorta, YS, bone marrow, placenta and embryonic head in mice (*25–29*). In human AGM, definitive HSPCs have recently been shown to emerge from *IL33*^+^*ALDH1A1*^+^ arterial endothelial cells (AEC) via *KCNK17*^+^*ALDH1A1*^+^ HE (*12*). Dissecting YS endothelial cell (EC) states in greater detail, the broad category of PVLAP^+^ ECs included AEC and HE, while LYVE1^+^ ECs encompassed sinusoidal, immature and VWF-expressing ECs (**Fig. 4A, S7A-B, Table S4-5**). HE was a transient feature of early YS (**Fig. 4A**). Along inferred trajectories, YS HSPCs appeared to arise from AEC via HE as in AGM (*11*), sequentially upregulating expected genes such as *ALDH1A1* (*30*) (**Fig. 4B**). The same EC intermediate states and transition points could be identified in both iPSC culture systems (**Fig. 4B, S7C**). In keeping with their more recent endothelial origin, we found that YS HSPCs and AGM HSPCs, but not embryonic liver or FBM HSPCs retained an EC gene signature characterised by the expression of *KDR, CDH5, ESAM* and *PLVAP* (**Fig. 4C**)

**Fig. 4:**
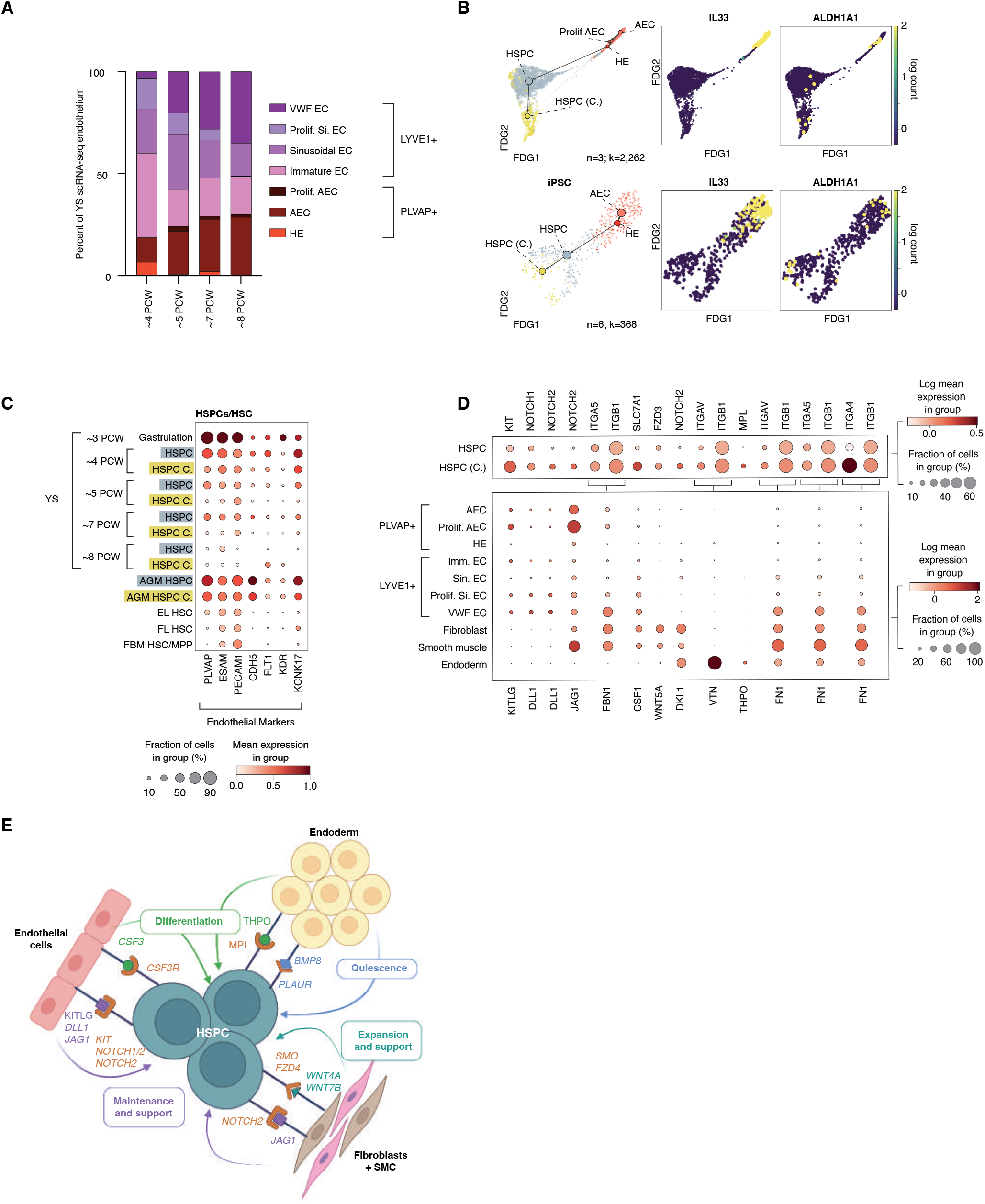
The lifespan of YS HSPCs. **(A)** Barplot showing the relative proportion of YS endothelial cell subsets grouped by PCW. **(B)** FDG overlaid with PAGA showing trajectory of HE transition to HSPC in YS scRNA-seq data (n=3; CS10, 11 and 14; k=2,262) (top) and iPSC-derived HSPC scRNA-seq data (n=7, k=437) (*21*) (bottom), with feature plots of key genes (*IL33, ALDH1A1*) involved in endothelial to hemogenic transition (**Table S5**). **(C)** Dot plot showing the expression level (colour scale) and percent expression (dot size) of genes associated with endothelial cells in HSPCs from YS (including gastrulation) scRNA-seq, AGM (*11*), matched EL (embryonic liver), FL (fetal liver) (*9*) and fetal bone marrow BM (*32*). **(D)** Heatmap showing relative mean expression z-scores of curated and statistically significant (p<0.05) CellphoneDB putative receptor ligand interactions between YS endothelial subsets vs HSPC across gestation. Growth factor, TGF beta and NOTCH ligand receptor -related gene interactions have been highlighted. **(E)** Schematic of selected and statistically significant (p<0.05) CellphoneDB putative receptor ligand interactions between YS HSPC vs endoderm, fibroblasts (Fib), smooth muscle cells (SMC) and endothelial cells (EC) in scRNA-seq data. Receptors and ligands in italics significantly decrease CS17-12 (6-8PCW). See CellPhoneDB methods and **Table S23-24**.

Receptor-ligand interactions capable of supporting YS HSPC expansion and maintenance were predicted using CellphoneDB (*31*). We identified ECs, fibroblasts, smooth muscle cells and endoderm as likely interacting partners (**Fig. 4D-E, Table S23**). ECs were predicted to maintain and support the HSPC pool through production of stem cell factor (KITLG) and NOTCH 1/2 ligands DLL1 and JAG1 (*32, 33*) (**Fig. 4D**). ECs potentially expand the HSPC pool via FN1 while fibroblasts and smooth muscle cells contribute via CSF1 and WNT5A (*34–36*) (**Fig. 4D**). WNT5A may also regulate lineage specification in YS HSPC as non-canonical WNT5A signalling promotes myelopoiesis at the expense of B lymphopoiesis in mouse (*37*). Endoderm is predicted to interact with HSPCs *via* VTN and THPO, reported to promote haematopoiesis and long-term-HSC-like quiescence, respectively (*38, 39*) (**Fig. 4D**).

Examining change in interactions over time, the key finding was diminishing interactions between CS17-CS23 (4-8PCW) including loss of cytokine and growth factor support, loss of TFGβ, WNT and NOTCH2 signals by all stromal fractions (**Fig. 4D-E, S7D, Table S24**). Many of these diminishing stromal ligands were still expressed in age-matched liver and AGM stromal cells (**Fig. S7E, Table S24**). Adhesive interactions in YS were also predicted to be significantly modulated (**Fig. S7F, Table S24**). While aged-matched liver provided opportunities for adhesion with stromal cells, AGM did not (**Fig. S7F, Table S24**). Interactions gained between CS17 and CS23 included endoderm-derived IL13 signalling to the *TMEM219*-encoded receptor implicated in apoptosis-induction (**Fig. 4D**). While limited conclusions can be made from studying cells that passed quality control for cell viability, we did observe upregulation in pro-apoptotic gene scores in late stage YS HSPCs, both primitive and definitive (**Fig. S7G**).

Despite marked change in the stromal environment of later stage YS, the proportion of HSPC to cycling HSPC remained stable (**Fig. S5F**). Differential lineage priming analysis revealed that very few HSPCs and mostly terminally differentiated cell states remained at CS22-23/8PCW YS (**Fig. S6C**) Together these observations are in keeping with an early burst of primitive HSPC production from transient YS HE, a later influx of definitive HSPCs derived from AGM, a loss of stromal support between 6-8 PCW, resulting in apoptosis and depletion of remaining HSPCs by terminal differentiation.

### An accelerated route to macrophage production in YS and iPSC culture

While YS progenitors are restricted to a short time-window in early gestation, mouse models suggest that they contribute to long-lived macrophage populations in some tissues (*40*). In our YS data, non-progenitor myeloid cells (lacking *AZU1, PRTN3* and *MPO*) included *HMGB2, LYZ* and *LSP1*-expressing promonocytes, pre-macrophages (expression profile below) and *C1QA/B/C* and *MRC* macrophages (**Fig. S8A**). Monocytes were observed only after liver-bud formation and AGM-derived haematopoiesis at CS14 (~5PCW) (**Fig. 5A, Fig. S8B).** A high probability of class prediction between YS and embryonic liver monocytes was noted (**Fig. 5B, Table S12**). A sub-population of YS monocytes (Monocyte2) expressed *ICAM3, SELL,* and *PLAC8* (adhesion molecules expressed by embryonic liver but not YS HSPCs), in keeping with liver-derived monocytes migrating to YS (**Fig. 5C)**. However, sequential waves of monopoiesis occurring within the YS cannot be excluded. YS CITE-seq data confirmed differentiatial expression of CD62L (*SELL*) and CD14 (*CD14*) on Monocyte2 compared to Monocyte1 and identified additional discriminatory markers; e.g., CD15, CD43 for Monocyte1 and CD9, CD35 for Monocyte2 (**Fig. S8C, Table S25**).

**Fig. 5:**
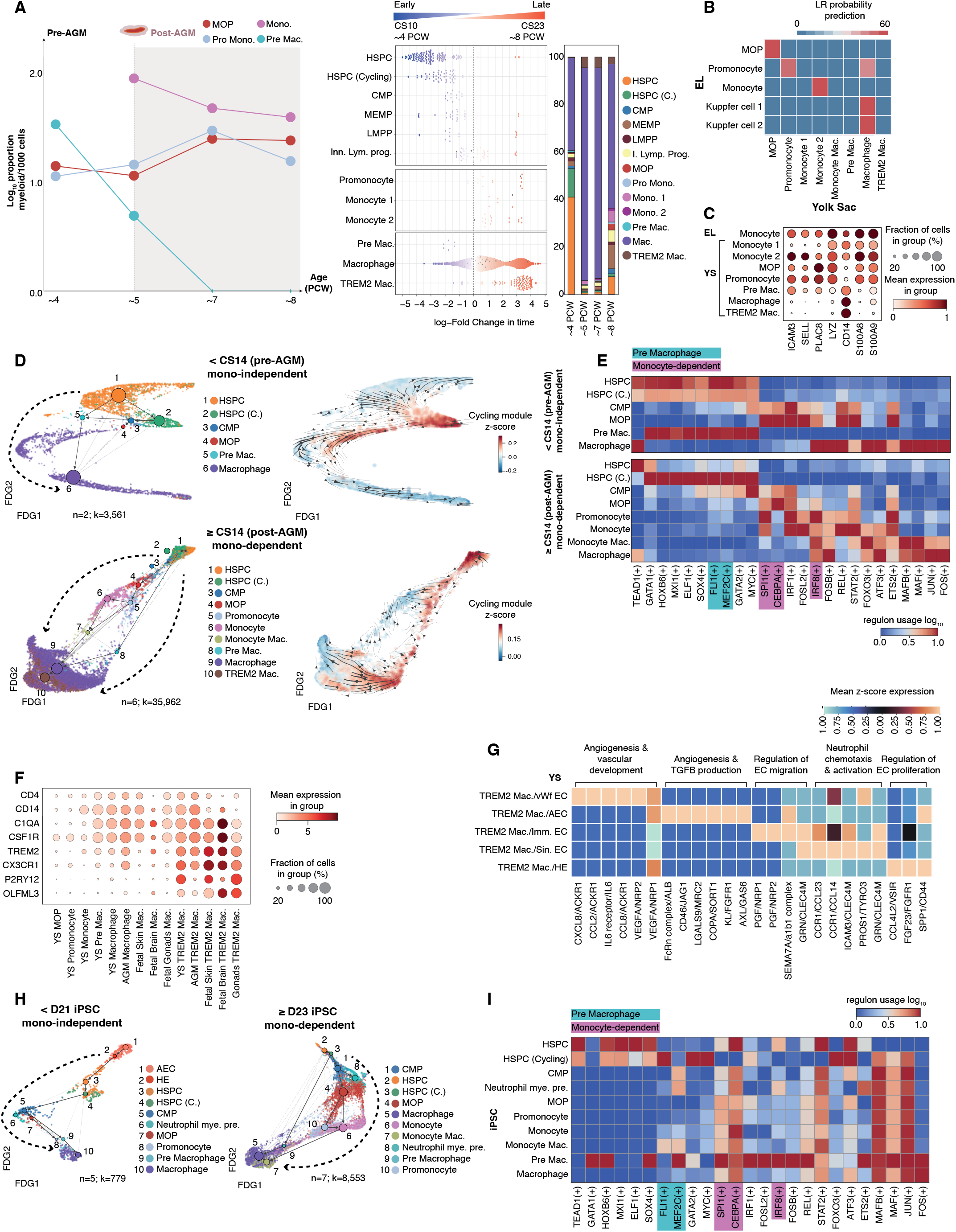
Accelerated macrophage production in YS and iPSC culture. **(A)** Left: Line graph showing the proportion of monocyte progenitors (MOP), promonocytes (pro mono), monocytes (mono) and pre macrophage (pre mac) in yolk sac (YS) over gestational age. Dashed line indicates the stage before (left) and after (right) AGM involvement. Middle: Milo beeswarm plot showing differential abundance of YS scRNA-seq myeloid neighbourhoods across time, where blue neighbourhoods are significantly enriched (SpatialFDR<0.1,logFC<0) early in gestation (CS10), red neighbourhoods are enriched later (CS23) (SpatialFDR<0.1,logFC>0) and colour intensity denotes degree of significance. HSPC= haematopoietic stem and progenitor cells, CMP= common myeloid progenitors, MEMP= mast, erythroid and megakaryocyte progenitors, LMPP= lymphoid-primed multipotent progenitors, inn. lymph. prog.= innate lymphoid progenitor, MOP= monocyte progenitor, mono= monocyte, mac=macrophage Prog.= progenitor, pre=, precursor, MK= megakaryocyte (**Table S19, 4**). Right: Bar graph showing the proportional representation of myeloid cell states in YS scRNA-seq data grouped by gestational age in PCW. **(B)** Median logistic regression class prediction probabilities for a model trained on YS scRNA-seq myeloid cell states (x-axis) projected onto equivalent cell states in matched embryonic liver (EL) scRNA-seq (y-axis)(**Table S12**). **(C)** Dot plot showing the expression level (colour scale) and percent expression (dot size) of DEGs and known monocyte markers in YS myeloid cell states shown in **b** compared to matched EL monocytes (standard_scale=‘var’) (**Table S28**). **(D)** Left: FDG overlaid with PAGA showing monocyte-independent trajectory from YS scRNA-seq HSPC to macrophage prior to CS14 (pre-AGM; n=2; k=3,561; top) and monocyte-dependent trajectory to macrophage after CS14 (post-AGM; n=6; k=35,962; bottom). Left: Coloured by cell state. HSPC, haematopoietic stem cell; CMP, common myeloid progenitor; MOP, monocyte progenitor (**Table S5**). Right: CellRank state transition matrix inferred arrows projected onto FDG indicate the trend of trajectory, and colour shows z-score enrichment in cycling module (GO:0007049) genes. **(E)** Matrix heatmap showing the pySCENIC-derived regulons associated with the YS macrophage trajectories as shown in **d** for monocyte-independent, and monocyte-independent routes of differentiation. **(F)** Dot plot showing the level (colour scale) and percent expression (dot size) of macrophage and microglia markers within the YS scRNA-seq monocyte and macrophage lineage in comparison to microglia, microglia-like and macrophage cell states in AGM (*11*), fetal skin (*45*), fetal gonads (*46*) and fetal brain (*44*). Fetal brain, skin and gonad cell states annotated in-house using LR output provided in **Fig. S8E.** Each dataset was scaled independently to a max_value = 10 then combined for plotting (**Table S18**). nocyte-dependent trajectories in the YS scRNA-seq dataset (including gastrulation). **(G)** Heatmap visualisation of CellphoneDB predicted interactions between TREM2^+^ microglia-like and endothelial cell states in the YS scRNA-seq (**Table S23**). Colour scale represents z-scored mean expression values of each gene pair. **(H)** FDG overlaid with PAGA showing monocyte-independent trajectory from iPSC scRNA-seq (*21*) AEC to macrophage prior to D21 (n=5; k=779; left) and monocyte-dependent trajectory to macrophage after D21 (n=7; k=8,553; right). Arrows indicate the trend of trajectory. Coloured by cell state (**Table S7, S5**). **(I)** Matrix heatmap showing the pySCENIC-derived regulons associated with the iPSC macrophage trajectories as shown in **i** for the iPSC scRNA-seq dataset (*21*).

The YS pre-macrophage uniquely expressed high levels of *PTGS2, MSL1* and *SPIA1,* as well as expressing progenitor genes (*SPINK2, CD34, SMIM24*), macrophage genes (*C1QA, MRC1*), and *CD52* which is typically associated with monocytes (**Fig. S8A**). This YS pre-macrophage rapidly declined by 5PCW and had no equivalent in embryonic liver (**Fig. 5A-B**), which led us to investigate putative macrophage differentiation trajectories in YS. FDG and partition-based graph abstraction (PAGA) visualisation revealed a direct, monocyte-independent trajectory to YS macrophages prior to CS14 (**Fig. 5D)**. In this pre-AGM trajectory, a transition from cycling HSPC to pre-macrophages, then macrophages (**Fig. 5D**) supports our predictions that primitive pre-AGM HSPCs exhibit myeloid bias (**Fig. 3G**). After CS14, a clear differentiation trajectory from cycling HSPC to monocytes and monocyte-macrophages was seen, but there were few transitional cells connecting to macrophages (**Fig. 5D)**. 15.33% of this macrophage pool was proliferating and CellRank RNA state transition analysis was in keeping with active self-renewal (**Fig. S8D**).

To explore how the two YS macrophage differentiation pathways are regulated, we examined differential transcription factor (TF) usage with PySCENIC (**Fig. 5E, Table S26**). YS premacrophages were predicted to use a group of TFs, including FLI1 and MEF2C, that have been reported in the differentiation of multiple lineages (*41, 42*), whereas the monocyte-dependent route (CMPs, MOP, promonocytes and monocytes) relied on recognised myeloid transcription factors such as SPI1, CEBPA and IRF8 (**Fig. 5E**).

TREM2^+^ macrophages (*43*), which had a high probability of correlation with fetal brain microglia (*44*), fetal skin TREM2^+^ (*45*) and fetal testes TREM2^+^ macrophages (*46*), were observed in YS only after CS14 (**Fig. 5A, 5D, 5F, S8E, Table S18**). By FDG, PAGA and CellRank RNA state transition analysis, TREM2^+^ macrophages were closely connected to the self-renewing macrophage population (**Fig. 5D, S8D**). *In situ*, YS TREM2^+^ macrophages were adjacent to the mesothelium, in a region enriched by EC (**Fig. S8F**). CellphoneDB predicted interactions between TREM2^+^ macrophages and VWF^+^ EC, via CXCL8 and NRP1, both of which are involved in angiogenic pathways (*47, 48*) (**Fig. 5G, Table S23**). TREM2^+^ macrophages also expressed the purinergic receptor P2RY12 that supports trafficking towards ATP/ADP-expressing ECs, as reported in mouse CNS (*49*) (**Fig. 5F**).

In a recent dissection of mouse macrophage heterogeneity across tissues and time, TLF^+^ macrophages (emerging from YS progenitors both directly and via a fetal monocyte intermediate) contributed to long-lived self-renewing tissue macrophage populations, while CCR2^+^ and MHC-II^hi^ macrophage pools received continual input from monocytes (*50*). To explore the contribution of YS-derived macrophages in prenatal human tissues, we interrogated a human pan-fetal immune cell atlas (*45*) to explore the proportion of cells representing each macrophage pool over gestational time (**Fig. S8G**). TLF^+^ cells first emerged in YS, and were subsequently found in both liver and skin. The proportion of TLF^+^ macrophages decreased over gestational time in all tissues, while the proportion of MHC-II^hi^ macrophages increased, particularly after 10PCW. While our data cannot discern whether TLF^+^ macrophages are upregulating an MHC-II^hi^ transcriptional programme or whether TLF^+^ macrophages are being replaced by a second wave of monocyte-derived macrophages, we can conclude that the TLF^+^ signature, temporally consistent with a YS origin, does not persist in humans to the extent observed in mouse (*50*). TLF^+^ macrophages in fetal liver were found within proliferating Kupffer cells (‘Kupffer 1’ in Popescu *et al* (*9*)). Liver and YS CITE-seq data confirmed protein expression of CD206 and CD163 in TLF^+^ macrophages and further identified CD28, CD49a, TSLPR and CD144 as positive identification markers that are minimally expressed in remaining Kupffer cells (**Fig. S8H, Table S27)**.

As macrophage differentiation from iPSCs permits high-resolution sampling over time, we sought to establish whether the macrophage subsets and developmental pathways we observed were recapitulated *in vitro*. To this end, we integrated our YS gene expression data with scRNA-seq data from iPSC-derived macrophage differentiation (n=19; k=50,512) by Alsinet et al (*21*) after refining the annotations of iPSC-derived cell-states (see Methods) (**Fig. S9A-D, Table S14, S5**). Non-adherent, CD14-expressing cells appearing after week 2 of differentiation expressed *C1QA, C1QB* and *APOC1* in keeping with a macrophage identity, while *S100A8/9, FCN1, CD52* and C*D14*-expressing monocytes only emerged after week 3 (**Fig. S9A-D**). Prior to monocyte emergence, a monocyte-independent macrophage differentiation trajectory was observed, consistent with the observations by Alsinet et al (*21*) (**Fig. 5H, Fig. S9A-D**). TF regulatory profiles of iPSC-derived macrophage differentiation were consistent with both pre-macrophage and monocyte-dependent *in vivo* TF profiles, including usage of *MEF2C, SPI1, CEBPA* and *IRF8* in iPSC-derived pre-macrophages (**Fig. 5I, Table 26**). However, neither system recapitulated TREM2^+^ macrophages suggesting that stromal cells, specifically ECs, may be required to acquire the TREM2^+^ molecular profile.

## Discussion

Using state-of-the-art single cell multiomic and imaging technologies we delineate the dynamic composition and functions of human YS *in vivo* from the 3rd post conception week (PCW), when the three embryonic germ layers form, to the 8th PCW when the majority of organ structures are already established (*22*). We detail how the YS endoderm shares metabolic and biosynthetic functions with liver and erythropoiesis-stimulating functions with liver and kidney. In part, this shared functionality may relate to a common role in creating a niche for haematopoiesis (*51*). Unlike in mice, where primitive erythroid progenitors mature in the YS (prior to circulation being established) but erythromyeloid progenitors can exit the YS and mature in the fetal liver, we show that active differentiation of erythroid and macrophage cells occurs for several weeks in human YS, prior to liver handover. The multi-organ functions, including extended haematopoiesis, of human YS may be an evolutionary adaptation to the longer gestation in humans. While an earlier study in humans, based on colony assays, suggested that the transition from YS to liver occurred at 5PCW, no YS samples were studied after this point (*52*).

The developmental window investigated here encompasses haematopoiesis from HSPCs arising both within the YS and within the embryo proper. We reconstruct YS HSPCs emergence from a temporally-restricted HE, featuring similar transition states and molecular regulation to AGM HSPCs. By gastrulation (CS7; ~3PCW), YS HSPCs already differentiate into primitive erythroid, MK and myeloid lineages. Building on a recent compilation of gene scorecards that characterise primitive and definitive HSPCs (*11*), we were able to parse the two fractions and document transition to definitive HSPC-dominance after CS14 (~5PCW). This separation also allowed us to identify a primitive HSPC bias towards myeloid, erythroid and megakaryocyte lineages and a definitive HSPCs bias towards megakaryocyte and lymphoid lineages. Both primitive and definitive HSPCs in the YS became more quiescent and upregulated apoptosis-related genes between CS17 and CS23 (~6-8PCW). Stromal cell ligands predicted to support HSPCs were markedly disrupted during this time, suggesting that the barriers to the survival of YS HSPCS may be extrinsic.

Primitive HSPCs uniquely employ an accelerated route to macrophage production independent of monocytes. The monocyte-dependent route may provide more tunable macrophage production via circulating innate immune cells to facilitate macrophage regeneration in response to tissue damage, inflammation or infection. While both primitive and definitive HSPCs, ‘accelerated’ and monocyte-dependent macrophages were recapitulated during *in vitro* differentiation of iPSCs, TREM2^+^ macrophages were not. TREM2^+^ macrophages, which are transcriptionally aligned with brain microglia, fetal skin, testes and AGM TREM2^+^ macrophages, were predicted to interact with endothelial cells, potentially supporting angiogenesis as described in mouse brain (*44*). Benchmarking of *in vitro* cultures against *in vivo* cell states and trajectories can facilitate more faithful replication of early blood and immune cells.

There is a growing appreciation of the potentially life-long consequences of early developmental processes. Our study illuminates a previously obscure phase of human development, where vital organismal functions are delivered by a transient extraembryonic organ employing non-canonical cellular differentiation paths. It will be fascinating to explore how these processes may impact on tissue homeostasis and disease across the human lifespan.

## Acknowledgements

For the purpose of Open Access, the author has applied a CC BY public copyright licence to any Author Accepted Manuscript version arising from this submission. We thank the Newcastle University Flow Cytometry Core Facility; Sanger Institute Cellular Genetics IT; Newcastle upon Tyne NHS Trust NovoPath; Tamilvendhan Dhanaseelan of HDBR for assistance with human fetal tissue processing and cell freezing; Natalina Elliott for contributions toward CITE-seq panel design.

## Author contributions

Conceived and directed the study: MH, SAT, BG

Acquired HDBR fetal samples: SL, DH

Generated scRNA-seq datasets: JE, ES, IG

Generated CITE-seq datasets: ES, NKW, NM, RH, MSV

Performed light-sheet microscopy: YG, MI

Performed immunofluorescence: DD, MA

Performed immunohistochemistry: RC

Performed RNAscope: KK, LT

Performed mouse and human embryo imaging: SJK

Generated and interpreted iPSC data: CAl, RVT, VL

Performed CITE-seq data analysis and interpretation: AR, MQL, NKW

Performed scRNA-seq data analysis and interpretation: IG, SW, AR, KG, IIR, DMP, KP, JPar, SvD

Interpreted the single cell data: OB, LJ, SBa, LG, MMa, KM, JPal, SBe, EL, AC, IR, MdB, ED, CS

Wrangled the single cell data: SW, MMa, MQL, AR, NKW, IG

Led web portal development: DH, DBL

Prepared the manuscript: BO, MMi

Drafted the manuscript: LJ, SW, RB, IG, MH

Designed the manuscript figures: JE, RB, CAd

## Data and materials availability

All novel raw sequencing data from this study are made publicly available at ArrayExpress as FASTQs and count matrices as follows:

- i) Human embryonic liver and yolk sac 10x scRNA-seq (E-MTAB-10552)
- ii) Human embryonic yolk sac 10x scRNA-seq (E-MTAB-11673)
- iii) Human embryonic yolk sac Smart-seq2 scRNA-seq (E-MTAB-10888)
- iv) Human embryonic yolk sac CITE-seq (E-MTAB-11549)
- v) Human embryonic liver CITE-seq (E-MTAB-11618)
- vi) Human fetal liver CITE-seq (E-MTAB-11613)

Accessions for published data reused in this study are detailed comprehensively in **Table S6**. Processed single cell datasets and supplementary tables are available for interactive exploration and download as well as corresponding trained scVI and logistic regression models via our interactive web portal (https://developmental.cellatlas.io/yolk-sac; password: ys2022). Of note, data on portals are best used for rapid visualisation - for formal analysis it is recommended to follow our GitHub code.

All code for reproducibility and trained logistic regression models of analysis is available at https://github.com/haniffalab/FCA_yolkSac

## Supplementary Materials

Materials and Methods

Figs S1 to S9

Refs 59 to 76

Tables S1 to S32

Movies S1 to S3

