## Supplementary Materials and Figures for "Multi-organ functions of yolk sac during human early development"

**This PDF file includes:**

Materials and Methods
Figs. S1 to S9
Captions for Movies S1 to S3
Captions for Tables S1 to S32

**Other Supplementary Materials for this manuscript include the following:**

Movies S1 to S3
Tables S1 to S32

### Materials and Methods

Ethics and sample acquisition

Tissues were obtained from the MRC–Wellcome Trust-funded Human Developmental Biology Resource (HDBR; http://www.hdbr.org) with appropriate written consent and approval from the Newcastle and North Tyneside NHS Health Authority Joint Ethics Committee (18/NE/0290). HDBR is regulated by the UK Human Tissue Authority (HTA; www.hta.gov.uk) and operates in accordance with the relevant HTA Codes of Practice. Tissues used for light-sheet fluorescence microscopy were obtained through INSERM's HuDeCA Biobank and made available in accordance with the French bylaw (Good practice concerning the conservation, transformation and transportation of human tissue to be used therapeutically, published on December 29, 1998). Permission to use human tissues was obtained from the French agency for biomedical research (Agence de la Biomédecine, Saint-Denis La Plaine, France).

Embryos were staged using the Carnegie staging method (*59*). A piece of skin or chorionic villi tissue was collected from each sample to perform quantitative PCR karyotyping of sex chromosomes and autosomal chromosomes 13, 15, 16, 18, 21 and 22 for the most commonly seen chromosomal abnormalities. No abnormalities were detected.

Please see **Table S6** for information regarding external scRNA-seq datasets that have been incorporated in this study.

Processing samples for imaging and single cell sequencing

Samples were dissected the same day and tissues transported in PBS on ice and processed immediately (<1 hour after dissection). For FFPE, samples were immediately placed in 10% (w/v) formalin and processed for embedding by NovoPath, Newcastle upon Tyne NHS Trust. For RNAscope, samples were snap frozen in an isopentane bath in LN2 prior to embedding in OCT (CellPath). For single cell suspensions, tissue was transferred to a sterile 10 $mm^2$ tissue culture dish and cut in <1 $mm^3$ segments before being transferred to a sterile 50 mL conical tube. Tissue was digested with pre-warmed digestion media (1.6 mg $ml^{-1}$ collagenase type IV (Worthington) in RPMI (Sigma-Aldrich) supplemented with 10% (v/v) heat-inactivated fetal bovine serum (FBS; Gibco), 100 U $ml^{-1}$ penicillin (Sigma-Aldrich), 0.1 mg $ml^{-1}$ streptomycin (Sigma-Aldrich), and 2 mM L-glutamine (Sigma-Aldrich)) for 30 min at 37 °C with intermittent shaking. Digested tissue was passed through a 100 μm filter, and cells were collected by centrifugation (500g for 5 min at 4 °C). Cells were treated with 1× RBC lysis buffer (eBioscience) for 5 min at room temperature and washed once with Flow Buffer (PBS containing 5% (v/v) FBS and 2 mM EDTA) before counting. Processing for scRNA-seq was continued promptly on fresh cells, for other uses (including CITE-seq) cells were collected by centrifugation (500g for 5 min at 4 °C) and resuspended in 10% (v/v) DMSO in FBS for freezing. For light-sheet fluorescence microscopy (LSFM), tissues were fixed in 4% PFA, dissected, and gestational age estimated as previously described (*60*).

Processing of single cell suspension for scRNA-seq

Immediately following isolation and counting, cells were collected by centrifugation (500g for 5 min at 4°C) and resuspended in residual buffer. 3 μL CD45 BUV395 (clone: HI30, BD Biosciences) was added to the resuspended cells and incubated on ice in the dark for 30 min, washed with Flow Buffer and resuspended at ~ 10 x $10^6$ cells $mL^{-1}$. Immediately prior to sorting, cells were passed through a 35 μm filter (Falcon) and DAPI (Sigma-Aldrich) was added at a final concentration of 3 μM. Flow sorting was performed on a BD FACSAria Fusion instrument using DIVA v.8, and data were analysed using FlowJo (v.10.4.1, BD Biosciences). Cells were gated to exclude dead cells and doublets, and then isolated for scRNA-seq analysis (droplet-based 10x Genomics, or plate-based Smart-seq2) using a 100 µm nozzle. For droplet-based scRNA-seq, $CD45^+$ and $CD45^-$ cells were sorted into separate chilled FACS tubes coated with FBS and prefilled with 500 µl sterile PBS. For plate-based scRNA-seq, $CD45^-AF^+SSC^{++}$ cells single cells were index-sorted into 96-well LoBind plates (Eppendorf) containing 10 µl lysis buffer (TCL (Qiagen) + 1% (v/v) β-mercaptoethanol) per well.

Library preparation and sequencing of scRNA-seq and CITE-seq samples

For the droplet-based scRNA-seq experiments, FACS sorted cell suspensions were counted and loaded onto the 10x Genomics Chromium Controller to achieve a maximum yield of 10,000 cells per reaction. 5' V1 kits were used and sequencing libraries were generated according to the manufacturer's protocols. Libraries were sequenced using either an Illumina HiSeq 4000 or NovaSeq 6000 to generate at least 50,000 raw reads per cell.

For the plate-based scRNA-seq experiments, the frozen cell lysates were thawed on ice for 1 minute. Purified cDNA was generated and amplified using a modified Smart-seq 2 protocol described in Villani et al (*61*). Sequencing libraries were then generated using Illumina Nextera XT kits with v2 index sets A, B, C and D. 384 cells were pooled and were sequenced using a HiSeq 4000 to generate at least 1 million raw reads per cell.

For the CITE-seq experiments, frozen cells were thawed and added to RF-10 media that had been pre-warmed to 37°C. Cells were then counted and pooled. Fc blocking reagent (Biolegend) was added to the cell pools and left to incubate at room temperature for 10 minutes. 0.5 µL CD34 APC/Cy-7 (clone: 581, Biolegend) was then added to the Fc blocked cells and left to incubate in the dark and on ice for 10 minutes. During this incubation, the CITE-seq antibody cocktail (Biolegend; **see Table S29**) was centrifuged at 14,000g for 1 minute, Flow buffer was then added to reconstitute before incubating for 5 minutes at room temperature. The resuspended antibody cocktail was then centrifuged at 14,000g for a further 10 minutes before adding to the cells. The cells and CITE-seq antibody cocktail were then left to incubate for 30 minutes in the dark and on ice. After this time, the cells were washed twice with Flow buffer and resuspended in a final concentration of 50 $\mu g\, ml^{-1}$ 7 AAD (Thermo Fisher Scientific) in Flow buffer.

Live, single cells or live, single $CD34^-$ cells and live, single $CD34^+$ cells (for the CITE-seq experiments) were then FACS sorted into 500 µl of PBS in FACS tubes coated with FBS. Cells were then counted and submitted to the CRUK CI Genomics Core Facility for subsequent processing using 10x Genomics protocols and sequencing. Single cell gene expression and cell surface protein libraries were generated using Single cell 3' v3 kits according to the manufacturer's protocol. Libraries were sequenced using a NovaSeq 6000 to achieve a minimum of 20,000 reads per cell for gene expression and 5,000 reads per cell for cell-surface protein.

Alignment, quality control, filtering, and preprocessing of scRNA-seq and CITE-seq data

scRNA-seq expression data (including droplet-based and plate-based) were mapped with CellRanger (version 3.0.2) to a human reference genome (see **Table S1**), and low quality cells expressing <2000 reads, <500 genes and >20% mitochondrial reads were filtered out of the data. Genes expressed in fewer than 3 cells were also filtered out of the data.

For droplet-based scRNA-seq data, the following additional QC steps were performed. Scrublet (*62*) v0.2.3 was applied to each sequencing lane for doublet detection, and clusters with

>(Median+(1.48*MAD)) of the median cluster doublet detection score were removed (**Table S3, F**). Ambient RNA was removed with Cellbender (fpr = 0.01, epochs = 150) (*63*) v0.2.0. To determine likelihood of maternal contamination, data were pooled by donor and submitted to Souporcell v2.4.0 at genotype clusters k=1 and k=2 models to represent likelihood of no maternal contamination and possible maternal contamination respectively. The optimal model was identified *via* BIC (Bayesian Information Criterion), where we observed a smaller BIC index at k=2 in one donor (F37, Female, 5PCW). Cells from the F37 alternate genotype were identified as potential maternal contaminants, composed mainly of Monocytes (n=149), and Mono_Macs (n=25), and excluded from downstream analysis.

For CITE-seq data, mapping was performed with Cellranger (v4.0.0) (GRCh38-2020-A reference genome) and CITE-seq-Count (v1.4.3) for RNA and ADT respectively, and deconvoluted using Souporcell singularity image at https://github.com/wheaton5/souporcell. Low quality cells expressing <200 genes and >20% mitochondrial reads were filtered out of the data and doublets were removed by applying Scrublet v0.2.3 to each sequencing lane and then removing clusters with >(Median+(1.48*MAD)) of the median cluster doublet detection score. CITE-seq protein data underwent QC and preprocessing similar to our previous study (*54*), i.e., cells were first filtered to intersect barcodes with counterpart CITE-seq RNA data, then unmapped antibodies were filtered out and then protein cells were filtered for low quality by cells with <30 proteins and expressing >5000 reads (**Table S4, S9, S30**).

For scRNA-seq count matrix transformation, normalisation and preprocessing were carried out with the Scanpy workflow (*64*). We normalised raw gene counts using the *sc.pp.normalize_total* function (*target_sum = 10e4*) from Scanpy (v1.9.0) in python (v3.8.6) and performed *ln(x)+1* transformation. Expression values reported are normalised, log-transformed and scaled to variance of mean using the *sc.pp.scale* function independently for each analysis.

For CITE-seq data count matrix transformation, we performed DSB-normalisation (denoised and scaled by background) (*65*) and then applied a Gaussian Mixture Model (GMM) for background signal regression. A modified DSB-normalisation approach used in our previous study (*54*) was constructed as follows. For each CITE-seq lane, low quality/empty droplets were identified as droplets under the largest UMI peak which had a value < 1.96*standard deviations (std) of the mean UMI counts value (mu_UMI) per sample. Peak detection was conducted using the *scipy.signal.find_peaks* function. The number of peak detection bins were dynamically estimated as (3.322*log(X)), where X was the total number of droplets. The model iterated through a series of 20 prominence intervals (0-20) with widths (0-10) where peaks detected < (mu_UMI-(1.96*std)) were retained as empty droplet peaks. In cases where no empty droplet peaks were detected, the empty droplet threshold was taken to be < (mu_UMI-(1.96*std)). The estimated empty droplets matrix was then taken into downstream DSB normalisation in the same way as our previous study (*54*). To further account for background protein signal, we used a gaussian mixture model (GMM) to model the mixture of protein expression levels in different cell types. We used the *sklearn.mixture.GaussianMixture* module fit 20 models with increasing number of cell clusters k (from k=2 and k=21) to represent expression of each protein by cell. The optimal model was identified using the sum of BIC (Bayesian Information Criterion) ($BIC_i = 2L_i + k_i \log n$) and AIC (Akaike information criterion) ($AIC_i = 2L_i + 2k_i$) where k is the number of GMM cell protein expression clusters, n is the number of cells in the sample and L is the model log likelihood. The model with the smallest sum of BIC and AIC indexes was selected. The expression values of the cluster with lowest expression from each GMM model were vertically stacked and used as a background mean per protein. The background protein mean was subtracted from DSB normalised protein expression values and summed per cell to create a background score per cell. The *scanpy.pp.regress_out* function was then used to regress against the per-cell background score to produce background-signal regressed counts.

Integration and batch correction of scRNA-seq and CITE-seq datasets

For integration of our in-house YS scRNA-seq dataset with external datasets, CellRanger count was first reapplied for the CS10/CS11 and CS14 embryonic YS scRNA-seq data acquired from Wang et al and Mikkola et al, respectively (*11*, *53*) (**Table S1**). The following steps were then followed for the total integrated YS droplet-based scRNA-seq dataset. Highly variable gene (HVG) selection was performed using the *sc.pp.highly_variable_genes* function (min_mean=0.001, max_mean=10) for embedding by dispersion. Dimensionality reduction and batch correction for the was carried out using the scVI (*66*) as used in scvi-tools (*67*) (dropout_rate=0.2, n_layer=2) with biological replicate taken as the technical covariate. To ensure model performance was optimal for each independent analysis, scVI was benchmarked against the python implementation of Harmony (*68*) (*Harmonypy* v0.0.5) at various theta values between 1 and 20. kBET (*69*) and Silhouette scores (*sklearn.metric.sil_score*) were computed for each iteration between donor covariates and compared to the scVI integration. Based on this, YS and liver CITE-seq data were batch corrected using the *Harmonypy* approach. Trained scVI models are provided via our interactive web portal.

Clustering and annotation of scRNA-seq and CITE-seq data

Clustering of scRNA-seq and CITE-seq datasets were performed using the leiden algorithm (*70*) (*sc.tl.leiden*) with a resolution parameter of *res*=1.5 on a k-nearest neighbourhood graph with k=15 unless specified otherwise. In cases where datasets are compared probabilistically, or where new classifications have been made, an implementation of low-dimensional ElasticNet regression (EN) (described in the 'Logistic regression' section of the manuscript methods) was used to first classify individual cells where a model-specific decision threshold of 0.9 was used for classification tasks. Cells classified inherited labels from the model trained on YS scRNA-seq data. Clusters were then assigned classes if the majority projected label had a label count distribution of > (mean + (1*std)) of label counts per cluster. Resultant cell state classifications were further manually checked using differentially expressed genes using the *sc.tl.rank_genes_groups* function in Scanpy which performed a two-sided Wilcoxon rank-sum test for genes expressed in >25% of cells, with a log-transformed fold change cut-off of 0.25. All p-values were adjusted for multiple testing using the Benjamini–Hochberg method. Annotation of YS and liver CITE-seq data was performed by training an EN model using YS scRNA-seq data as reference. These labels were then distributed by majority voting onto leiden clusters derived from CITE-seq data **(Table S31, S10)**. The resultant cluster annotations were validated using the same markers identified in matched RNA data and underwent additional manual annotation where required.

Dimensional reduction and marker expression visualisation

For visualisation, the uniform manifold approximation (UMAP) algorithm was run using the *sc.tl.umap* function in Scanpy. Dot-plots and violin plots were produced in Scanpy and all gene expression values displayed were normalised, log-transformed, and scaled as described in the preprocessing section unless otherwise stated. Force directed graphs (FDGs) computed with the *sc.tl.draw_graphs* function in Scanpy using the Force Atlas2 parameter were used to infer trajectories. PAGA graph abstractions were computed on the k-nearest neighbour graphs and overlaid onto FDGs where nodes represented the centroid of each cell state cluster and the thickness of edges represented the similarity between cell states (**Table S5**).

Proportion line graphs were produced with Matplotlib for cells enriched in specific genes (e.g., *HBZ*) where enrichment was defined by (>0) average z-scored expression of each gene subtracted with the average expression of a randomly sampled set of 50 reference of genes at 25 bins using the *sc.tl.enrich* function in Scanpy. Proportions of enriched cells with z-score >0 in each cell type compartment (e.g., erythroid cells) were then graphed across timepoints to visualise differential enrichment of cells expressing the genes of interest.

<u>Differential abundance testing and FACS correction</u>

We tested for differential cell-state abundance across gestation using the Milo framework, correcting for CD45 positive and negative FACS isolation strategies using the technique employed in Suo *et al* 2022 (*45*), where we calculated a FACS isolation correction factor for each sample $s$ sorted with gate $i$ as ($fs = \log(p_i S/S_i)$) where $p_i$ is the true proportion of cells from gate $i$ and S represents the total number of cells from both gates. A KNN graph was then constructed from the remaining cells using the *milopy.core.make_nhoods* function (prop = 0.05). Neighbourhood by majority frequency of cell labels in each neighbourhood (>50%). The YS scRNA-seq data was then split into 5 age bins (3PCW, 4PCW, 5PCW, 7PCW and 8PCW) and cell counts were modelled as a negative binomial generalised linear model (NB-GLM) with Benjamini-Hochberg weighted correction as described in Suo *et al (45).* Significantly differentially abundant neighbourhoods were detected by SpatialFDR (<0.1, logFC <0) for early enriched neighbourhoods and SpatialFDR (<0.1, logFC >0) for late neighbourhoods. (**Table S19**).

<u>Clustered gene-set enrichment analysis</u>

We ranked conserved markers (p-value < 0.05) between the endoderm cell state in YS scRNA-seq data against hepatocytes in EL scRNA-seq and endoderm in the mouse gastrulation scRNA-seq data using the *FindConservedMarkers* function in Seurat (v3.1) with Bonferroni corrected FDR adjusted p-values. Markers were submitted for gene set enrichment ranking and analysis using the Enrichr tool as implemented in the *GSEApy* package to query the Gene Ontology (GO) Biological Process database (GO_BP_2022) (**Table S22**). Using the enrichrR package (v3.0) in R, enrichment was first computed by Fisher exact test for randomly sampled genes to derive a mean rank and standard deviation to estimate background for each ontological term accessed. A z-score for deviation of each term to its background rank was then used to rank output genesets. We derived statistical significance (Fisher exact test < 0.05, ranked by Z-score) for each gene set enrichment and performed Markov clustering (MCL) using the MCL (v1.0) package in R to derive network neighbourhoods based on geneset intersect. Gene set clusters were annotated using the AutoAnnotate function in the RCy3 (v2.16) package and clusters were ranked by the mean Z-score of all gene sets within each cluster and manually curated based on biological significance. We used the Cytoscape software (v3.9.1) to visualise clusters.

<u>Cell state predictions using probabilistic low-dimensional ElasticNet regression</u>

Label transfer class assignments and median probability of class correspondence between gene expression matrices in single cell datasets were carried out using a Logistic Regression

(LR) framework as applied in our previous work (*54*) using a similar workflow to the CellTypist tool (*71*).

Raw gene expression scRNA-seq datasets being compared were first concatenated, normalised and log-transformed as described in pre-processing. HVG selection was performed (min_mean=0.001, max_mean=10) for embedding by dispersion and dimensionality reduction carried out with PCA (k=100 components unless otherwise specified). Batch correction was carried out using Harmonypy with donor and dataset origin information taken as technical covariates. Harmony runs were iterated through theta=1:20 and resultant embeddings benchmarked using kBET and silhouette scores between technical covariates where a low kBET rejection rate and corresponding high silhouette score denoted the optimal theta parameter.

An ElasticNet regression (EN) LR model was built utilising the 'sklearn.linear_model.LogisticRegression' module in the sklearn package (v0.22). The model was trained using the sliced batch-corrected low-dimensional PCA representation of the training data with regularisation parameters tuned using the GridSearchCV function in *sklearn.* The test grid was designed with five l1_ratio intervals (0 , 0.2, 0.4, 0.6, 0.8 ,1) at 5 train-test splits and 3 repeats for cross-validation. The unweighted mean over the weighted MSEs of each test fold (the cross validated MSE) was used to determine the optimal model. To avoid situations where class imbalances contributed to low predictive probabilities of under-sampled classes, the SMOTE workflow from the *imblearn* package was utilised to resample populations > (mean+(1*std)) whilst preserving important features (*k_neighbours* = 15) prior to training.

The resultant model was used to predict the probability of correspondence between trained labels and pre-computed clusters in the target dataset. For dataset comparison tasks where pre-designated labels already existed in the target dataset, the median probability of training label assignment per pre-designated class was computed and visualised as a heatmap (**Table S11-18**).

For classification tasks, a model-specific decision threshold of 0.9 was used to determine predicted labels. Clusters were then assigned classes if the majority projected label had a label count distribution of > (mean + (1*std)) of label counts per cluster. Resultant cell state classifications were further manually checked using differentially expressed genes. Further assessment of the predicted cluster labels was carried out by computing the Adjusted Rand index and Mutual information scores from the modules 'sklearn.metrics.adjusted_rand_score' and 'sklearn.metrics.mutual_info_score' between the original cluster labels and predicted cluster labels in each dataset. This methodology was applied to classify and annotate several external datasets including the scRNA-seq human gastrulation data, the human AGM data, the human embryonic liver data and human fetal skin data, as well as the human YS and liver CITE-seq data.

An implementation of the EN workflow described above, in conjunction with the SAMap (Self-Assembling Manifold mapping) workflow (*72*) was used to classify and probabilistically compare cell states across the human YS scRNA-seq data and the mouse gastrulation YS data. A gene-gene sequence homology graph weighted by human and mouse protein sequence similarity was first constructed using the SAMAP command. Reciprocal BLAST mapping using the tblastx tool between the entire mouse and human transcriptomes for significant homology (E-value $<10^{-6}$) was supplied. The resultant SAM object returns *k*=300 species-stitched PC components for the top 3000 paired genes. These PC components were used to train the cross-species EN model as a classification task described above (**Table S13**).

Trained Logistic regression models are provided via our interactive web portal in '.sav' format and may be accessed using the Python package CellTypist (v.0.1.9)(*71*) for projection and prediction of external scRNAseq datasets.

Differential lineage priming and progenitor cell fate predictions

The CellRank package (v1.5.1) was used to define and rank fate probabilities of terminal state transitions across annotated haematopoietic lineages in the YS and iPSC scRNA-seq datasets. First order kinetics matrices were imputed for each dataset using the *pp.moments* function (n_pcs=20, n_neighbours=30) in the scVelo package (v0.2.4). A Cytotrace pseudotime for state-transitions across each dataset was then computed to direct graph-edges towards estimated neighbourhood regions of increasing differentiation using the Cytotrace kernel provided within the CellRank package. The resultant KNN and Cytotrace pseudotime were used to compute a probability transition matrix with the *compute _transition_matrix* command in Cytotrace. Neighbourhoods of cells representing terminal states of differentiation were identified using true Schur matrix eigen decomposition of the transition matrix *compute_schur* (n_components=20, method='brandts'), followed by the *compute_macrostates* (n_states=10) command in Cytotrace. The resultant terminally differentiated cell states were then manually selected if multiple terminal states were identified per lineage. Fate absorption probabilities were then computed across all cells terminating at each pre-specified terminal cellstate neighbourhood using the *compute_absoprtion_probabilities* command in CellRank. Fate probabilities were then presented as a circular plot using the *pl.circular_projection* with embedding proximity to terminal edges of the graph representing the fate-transition probability of a particular cell towards the pre-specified terminally differentiation state. HSPC progenitor population density was then computed by kernel density estimation (KDE) of a pre-computed UMAP highlighting relative probabilities of HSPC lineage priming (KDE calculated using the *tl.embedding.density* function in Scanpy).

pySCENIC for regulon analysis

The PySCENIC package (v0.9.19) was used to identify transcription factors and their target genes in the YS and iPSC scRNA-seq datasets. The ranking database (hg38 refseq-r80 500bp_up_and_100bp_down_tss.mc9nr.feather), motif annotation database (motifs-v9-nr.hgnc-m0.001-o0.0.tbl) and list of transcription factors (lambert2018.txt) were used. An adjacency matrix of transcription factors and their targets was generated. TF activity from the AUcell output was modelled along diffusion pseudotime rankings of each trajectory and used to train a Non-linear Generalised Additive Model (nlGAM) using the pyGAM.LinearGAM model to identify TF modules which significantly changed across each lineage pseudotime. A gridsearch of between 50 and 200 splines were calculated. Significantly changing TF regulons across pseudotime were classified with a p-value < 0.05 and reported in **Fig. 5E, 5I (Table S26)**.

Cell-cell interaction predictions using CellPhoneDB

To assign putative cell-cell interactions within the YS scRNA-seq dataset, we used CellPhoneDB (v2.1.2). Log-transformed, normalised and scaled gene expression values for all cell states were exported. CellPhoneDB was run using the statistical method using the receptor-ligand database (v2.0.0) with significance p-value cut-off of 0.05 and a result precision of 3dp (**Table S23-24**). Outputs were ranked by log-mean expression for interactions between cell types of interest in each analyses and plotted as a z-scored heatmap to show standard deviations from mean for each receptor-ligand pair.

Hiplex RNAscope

Human yolk sac tissue (8PCW) was frozen in optimal cutting temperature compound (Tissue-Tek). 12-plex smFISH was performed using the RNAscope HiPlex v2 assay (ACD, Bio-Techne) on 3 cryosections (10 µm) per manufacturer's instructions, using the standard pre-treatment for fresh frozen samples and permeabilized with Protease III, for 15 mins at room temperature. The imaging cycles, primary probes and label fluorophores were: *Cycle1_KLRB1_AlexaFluor488, Cycle1_CD1C_Dylight550, Cycle1_IL7R_Dylight650, Cycle1_SPINK2_AlexaFluor750, Cycle2_P2RY12_AlexaFluor488, Cycle2_TNFA_Dylight550, Cycle2_LGALS3_Dylight650, Cycle2_IL33_AlexaFluor750, Cycle3_PLVAP_AlexaFluor488, Cycle3_SPINK1_Dylight550, Cycle3_C1QA_Dylight650, Cycle3_ACTA2_AlexaFluor750, Cycle4_P2RY12_Opal570 and Cycle4_IBA1_Cy5.* Finally, the slides were counterstained with DAPI and coverslipped for imaging.

For protein validation, slides were fixed with 4% (w/v) PFA for 60 minutes at room temperatures, then washed and dehydrated in an ethanol gradient (50-100%) for 5 minutes each. Sections were treated with Protease III (ACD, Bio-Techne) for 15 minutes at room temperature, then washed with PBS prior to blocking in 10% (v/v) normal donkey serum containing 1% (w/v) Triton X-100 and 0.2% (w/v) gelatin for 60 minutes at room temperature. Primary antibody staining was done at 4°C overnight, then washed three times for 20 minutes each with Wash buffer (0.1% (w/v) Triton X-100 in PBS). Slides were then blocked with HRP Block (ACD, Bio-Techne) for 60 minutes at room temperature, and washed with ACD Wash Buffer (ACD, Bio-Techne) prior to addition of secondary antibody and incubated for 60 minutes at room temperature. Slides were washed three times for 20 minutes each with Wash buffer (0.1% (w/v) Triton X-100 in PBS), TSA-Opal570 added for 10 minutes at room temperature, then washed three times with ACD Wash Buffer. Finally, the slides were counterstained with DAPI and coverslipped for imaging.

Imaging was performed on a custom two camera spinning disk confocal microscope built around a Crest Optics X-light v3 module by Cairn Research, a scientific equipment manufacturer. The instrument was controlled using the Micro-Manager software (*73*). All imaging was done in spinning disk confocal mode using a 40x water immersion objective (NA 1.15, 180nm/pixel) with a 1.5 µm z-step.

RNAscope image analysis

Before each imaging experiment, a slide covered in a sparse layer of 0.5 µm Tetraspeck beads was also imaged in all channels. The bead images in all channels were then registered against the beads in the DAPI channel and their respective affine transforms were saved.

After imaging, each individual tile was z-projected with a maximum intensity projection, then the channels were transformed using the saved affine transforms. The projected, transformed tiles were then saved back to a temporary directory along with a bigstitcher-compatible XML file. The BigStitcher software (*74*) was then used to stitch the transformed tiles together and the final stitched image exported for further analysis.

All imaging cycles for a given tissue section were then registered in two steps. Firstly, we used feature registration algorithm implemented in Python via OpenCV-contrib library (version

4.3.0) (*75*) to compute an affine transformation of DAPI channel from round r>1 (moving image) with respect to DAPI channel from the first round r=1 (reference image). In particular, key points were detected using the FAST feature detector, whose surrounding areas were described using the DAISY feature descriptor, while the FLANN-based matcher was used to find correspondences between pairs of key points from reference and moving images and filter out unreliable points. Lastly, the remaining key points were processed using the RANSAC-based algorithm that aligns them and estimates affine transformation parameters with four degrees of freedom.

For the second registration step, a nonlinear registration algorithm based on Farneback optical-flow available in Python via OpenCV library was used to achieve more accurate registration by warping images locally. Specifically, local warping was computed using the anchor channel (mouse brain and lymph node), or DAPI channel (human brain), from round r>1 with respect to the corresponding channel of the first round. The computational pipeline implementing these registration steps was optimised so that it can be performed efficiently on large images and the corresponding code for feature registration is available at github.com/BayraktarLab/feature_reg, while the code for optical-flow registration at github.com/BayraktarLab/opt_flow_reg.

Immunohistochemistry

Formalin-fixed, paraffin-embedded blocks of yolk sacs aged 4-8PCW, embryonic livers aged 7-8PCW, and healthy adult livers were sectioned at 4 μm thickness onto APES-coated slides.

For Haematoxylin and Eosin staining, slides were dewaxed in xylene and rehydrated through graded ethanol, as previously published (*9*). Rehydrated slides were incubated for 5 minutes in Mayer's haematoxylin (Dako, Agilent), rinsed in tap water and then differentiated for 2 seconds in acid alcohol before washing in tap water followed by Scott's tap water substitute (Leica Biosystems). Sections were counterstained in triple eosin (Dako, Agilent) for 5 minutes before being rinsed in tap water, dehydrated through graded ethanol (70% to 99%) and then placed in xylene before mounting with DPX (Dako, Agilent).

For IHC, dewaxing, rehydration and staining was done using the Discovery Ultra auto Stainer and kits (Ventana, Roche). After rehydration, one drop of Inhibitor (Ventana, Roche) was added and incubated for 8 minutes before rinsing with Reaction Buffer (Ventana, Roche). One drop of primary antibody (**Table S32**) was added and incubated for 32 minutes prior to rinsing with Reaction Buffer. One drop of OMap anti-mouse (CK19) or anti-rabbit (ALB and alpha-1-antitrypsin) HRP (Ventana, Roche) was added and incubated for 16 minutes prior to rinsing with Reaction Buffer. One drop of hydrogen peroxide (Ventana, Roche) was applied to the slide, which was incubated for 4 minutes before the addition of one drop of DAB (Ventana, Roche) and an additional 8 minute incubation. The slide was washed with Reaction Buffer then one drop of Copper (Ventana, Roche) applied and incubated for 4 minutes prior to washing with Reaction Buffer. The slide was counterstained with one drop of hematoxylin II (Ventana, Roche) for 8 minutes, rinsed with Reaction Buffer and one drop of Bluing reagent (Dako, Agilent) added for 4 minutes. The slide was then rinsed with a Reaction buffer, before being dehydrated by hand through graded ethanol (70% to 99%), placed in xylene and mounted with DPX (Dako, Agilent).

Rabbit polyclonal anti-human alpha-1-fetoprotein (AFP; Agilent) staining was done by the Newcastle upon Tyne NHS diagnostic lab using a proprietary method.

For $CD34^{+}VEGFR2^{+}$ co-staining, dewaxing, rehydration and staining was done using the

Discovery Ultra auto Stainer and kits (Ventana, Roche). After rehydration, one drop of DISC inhibitor (Ventana, Roche) was added to the slide which was then incubated for 8 minutes prior to rinsing with Reaction Buffer. One drop of mouse monoclonal anti-human CD34 (clone QBEnd/10; Roche; **Table S32**) was added, incubated for 32 minutes then rinsed with Reaction Buffer. One drop of OMAP anti-mouse HRP (Ventana, Roche) was added and incubated for 16 minutes then rinsed with Reaction Buffer. Two drops of DISC purple (Ventana, Roche) were added, incubated for 4 minutes, then one drop of hydrogen peroxide (Ventana, Roche) added for a 24 minute incubation. The slide was washed with a Reaction Buffer then 2 drops of DISC inhibitor (Ventana, Roche) added. The slide was incubated for 20 minutes, rinsed with Reaction Buffer then one drop of rabbit mono-clonal anti-human VEGFR2 (clone: D5B1; Cell Signalling Technologies) added and incubated for 32 minutes. The slides were rinsed with Reaction buffer prior to the addition of one drop of UMap anti-rabbit AP (Ventana, Roche), incubated for 16 minutes then rinsed with Reaction Buffer. Three drops of yellow Buffer (Ventana, Roche) were applied, incubated for 4 minutes, then one drop of Disco Yellow (Ventana, Roche) applied and incubated for 40 minutes. The slide was rinsed with Reaction Buffer then counterstained with one drop of hematoxylin II (Ventana, Roche) for 8 minutes, rinsed with Reaction Buffer, and incubated for 4 minutes after the addition of one drop of Bluing reagent (Ventana, Roche). The slide was then rinsed with a Reaction buffer, before being dehydrated by hand through graded ethanol (70% to 99%), placed in xylene and mounted with DPX (Dako, Agilent).

For the Martius Scarlet eBlue (MSB) stain, slides were dewaxed in xylene and rehydrated through graded ethanol as previously published (*9*). Rehydrated slides were placed in Bouins' fixative (Atom Scientific) for 1 hour at 60°C, washed in running water, incubated in Weigert's solution (Atom Scientific) for 10 minutes and washed in water. Slides were differentiated in 0.9% ethanol for 1-2 seconds before rinsing in tap water followed by Scott's tap water substitute (Leica Biosystems), distilled water and finally 95% ethanol. Slides were then incubated stepwise in Martius yellow (3 minutes) (Atom Scientific), Brilliant crystal scarlet (6 minutes) (Atom Scientific) then 50% (v/v) Methyl blue (2 minutes) (Atom Scientific), washing with distilled water between each stain before a final tap wash and rapid dehydration (2-3 mins) through graded ethanol (70% to 99%) and then placed in xylene before mounting with DPX (Dako, Agilent).

All slides were imaged at 20x magnification on a NanoZoomer S360 (Hamamatsu) digital slide scanner. MSB stained images were deconvolved into respective Martius yellow, crystal scarlet and methyl blue channels using the Colour Deconvolution plugin (v1.8) (Masson Trichrome) in FIJI with thresholds set using the Otsu method. Pseudo-colours for each deconvolved channel were then assigned as in **Fig. 2C.**

<u>ASGR1 and CD34 Immunofluorescence microscopy</u>

Yolk sac sections were baked onto slides for 2 hours at 60°C before being dewaxed in xylene and rehydrated through graded ethanol as previously published (*9*). Slides were washed with distilled water then placed in a pressure cooker with boiling citrate buffer pH 6 (10mM citric acid (Sigma), 0.05% v/v Tween 20 (Sigma) in DI water) for 2 minutes for antigen retrieval. Slides were then washed for 3 minutes with distilled water followed by 3 minutes in PBS (Sigma). Sections were blocked with 20% (v/v) goat serum (R&D Systems) for 45 minutes at room temperature. Primary antibodies were made up in blocking solution (see **Table S32**), added to the sections and incubated for 1 hour at room temperature. Slides were washed twice for 3 minutes each in wash buffer (0.1% (w/v) Triton X (Sigma) in PBS) then twice for 3 minutes each in PBS. Secondary antibodies (see **Table S32**) were made up in blocking solution, added to section and incubated for 2 hours at room temperature. The wash step was repeated then 300nM DAPI (Sigma) added, incubated for 5 minutes, washed with PBS and slides mounted with ProLong™ Diamond Antifade (Thermofisher). Slides were imaged on a Zeiss Axioimager with Zeiss ZEN pro software.

SMA and LYVE1/CD34 Immunofluorescence microscopy

PFA-fixed yolk sacs were cryoprotected with sucrose 10%, embedded in gelatin-sucrose solution (7.5% w/v gelatin (VWR 24350.262), 10% w/v sucrose (VWR27478.296), in 0.12M PBS), frozen at -50°C, then sectioned at 14um. Slides were stored at -80°C until use, dried for 30 minutes, then blocked with PBS Gelatin Triton (0.2% w/v gelatin, 0.25% w/v Triton X-100 (Sigma-Aldrich) in PBS) for 1 hour. Primary antibodies were made up in blocking solution (**Table S32**), added to the sections, and incubated overnight. Slides washed with PBS 3 times at 10 minute intervals. Secondary antibodies were made up in blocking solution and added to sections to incubate for 2 hours (**Table S32**). Hoechst 33258 (Sigma-Aldrich) was added to the secondary antibody solution. Sections were washed with PBS 3 times at 10 minute intervals, and coverslips were mounted with Mowiol (Calbiochem). Sections were imaged at 20x magnification on Leica DM6000 widefield microscope with MetaMorph software. Brightness and contrast were adjusted and a scale bar was added with FIJI (*76*).

Light-sheet fluorescence microscopy

Embryonic and fetal yolk sacs were dissected with their connecting stalk whenever possible from morphologically normal specimens collected from 5 to 9 post conceptional weeks (PCW). Candidate antibodies were screened by immunofluorescence on cryosections obtained from OCT-embedded specimens as previously described (*9*, *60*). Routine light-sheet immunofluorescence microscopy (LSFM) was then performed on floating whole-mount yolk sacs as previously described, with primary antibody incubation reduced to ten days and secondary reduced to two days, both at 37°C to preserve tissue integrity. Antibody and other reagents including nuclear marker TO-PRO-3 Iodide are specified in (**Table S32**). Yolk sacs were embedded in 1.5% agarose blocks prior to solvent-based clearing as previously described (*60*). Remarkably, yolk sacs retained their spherical shape throughout the procedure. Imaging was performed as previously described in dibenzyl ether with a Miltenyi Biotec Ultramicroscope Blaze (sCMOS camera 5.5MP controlled by Imspector Pro 7.3.2 acquisition software), which generates light sheets at excitation wavelengths of 488, 561, 640, and 785 nm. Objective lenses of 4x magnification (MI Plan 4x NA0.35) and 12x magnification (MI Plan NA 0.53) were used. Imaris (v9.8, BitPlane) was used for image conversion, processing, and video production. Photoshop (Adobe) was used to create panels. Blender 3.0 was used to edit videos and add text. All raw image data are available on request (AC, MH).

Statistics and reproducibility

The number of cells from each cell type in each de novo single cell dataset provided in this manuscript are provided in **Table S4.**

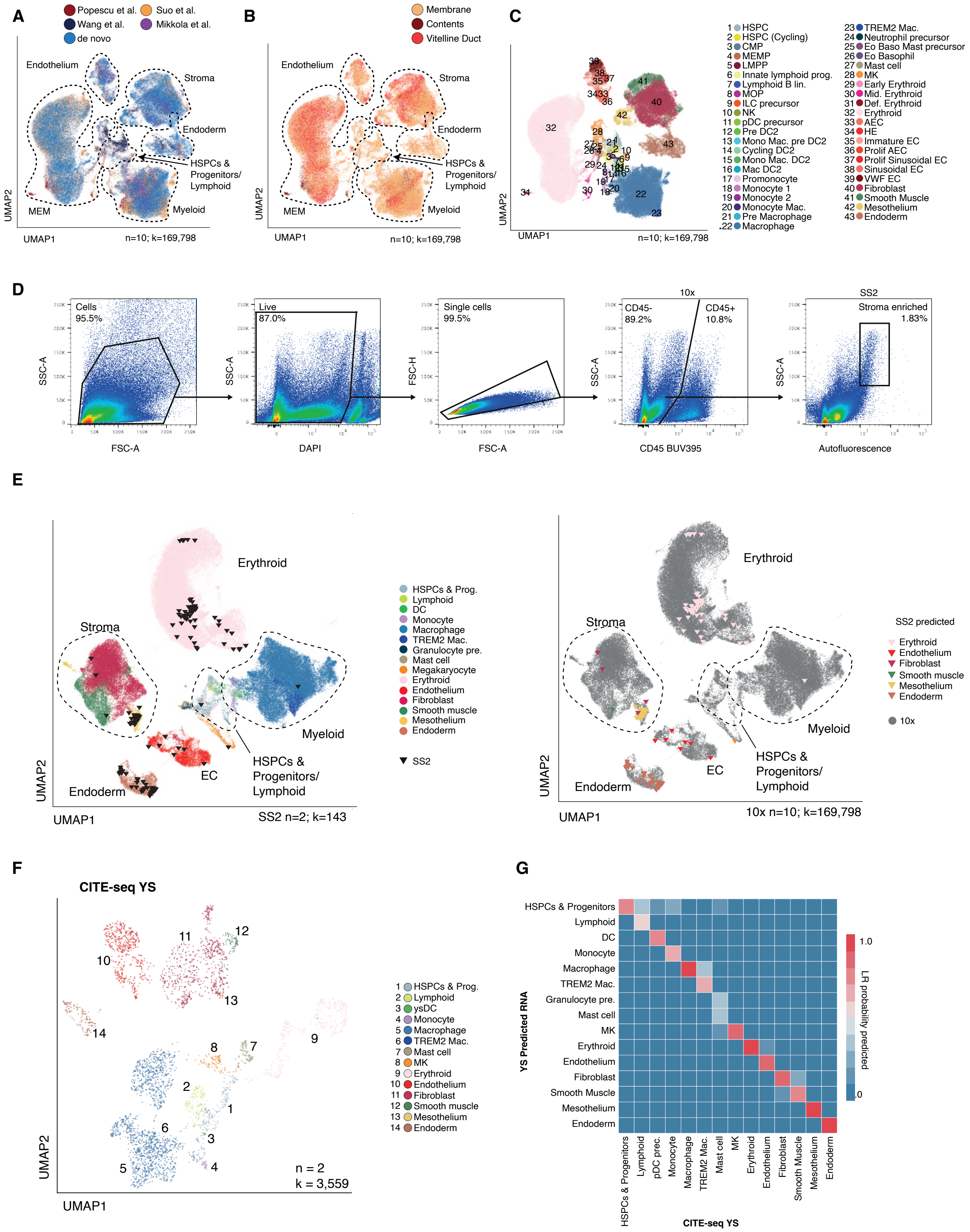

A
Popescu et al.
Suo et al.
Wang et al.
Mikkola et al.
de novo
Endothelium
Stroma
Endoderm
HSPCs & Progenitors/ Lymphoid
MEM
Myeloid
UMAP2
UMAP1
n=10; k=169,798
B
Membrane
Contents
Vitelline Duct
C
1 HSPC
2 HSPC (Cycling)
3 CMP
4 MEMP
5 LMPP
6 Innate lymphoid prog.
7 Lymphoid B lin.
8 MOP
9 ILC precursor
10 NK
11 pDC precursor
12 Pre DC2
13 Mono Mac. pre DC2
14 Cycling DC2
15 Mono Mac. DC2
16 Mac DC2
17 Promonocyte
18 Monocyte 1
19 Monocyte 2
20 Monocyte Mac.
21 Pre Macrophage
22 Macrophage
23 TREM2 Mac.
24 Neutrophil precursor
25 Eo Baso Mast precursor
26 Eo Basophil
27 Mast cell
28 MK
29 Early Erythroid
30 Mid. Erythroid
31 Def. Erythroid
32 Erythroid
33 AEC
34 HE
35 Immature EC
36 Prolif AEC
37 Prolif Sinusoidal EC
38 Sinusoidal EC
39 VWF EC
40 Fibroblast
41 Smooth Muscle
42 Mesothelium
43 Endoderm
D
Cells 95.5%
Live 87.0%
Single cells 99.5%
10x
CD45- 89.2%
CD45+ 10.8%
SS2
Stroma enriched 1.83%
SSC-A
SSC-H
FSC-A
DAPI
CD45 BUV395
Autofluorescence
E
Erythroid
Stroma
Myeloid
EC
Endoderm
HSPCs & Progenitors/ Lymphoid
HSPCs & Prog.
Lymphoid
DC
Monocyte
Macrophage
TREM2 Mac.
Granulocyte pre.
Mast cell
Megakaryocyte
Erythroid
Endothelium
Fibroblast
Smooth muscle
Mesothelium
Endoderm
SS2
SS2 n=2; k=143
SS2 predicted
10x
10x n=10; k=169,798
F
CITE-seq YS
1 HSPCs & Prog.
2 Lymphoid
3 ysDC
4 Monocyte
5 Macrophage
6 TREM2 Mac.
7 Mast cell
8 MK
9 Erythroid
10 Endothelium
11 Fibroblast
12 Smooth muscle
13 Mesothelium
14 Endoderm
n = 2
k = 3,559
G
HSPCs & Progenitors
Lymphoid
DC
Monocyte
Macrophage
TREM2 Mac.
Granulocyte pre.
Mast cell
MK
Erythroid
Endothelium
Fibroblast
Smooth Muscle
Mesothelium
Endoderm
YS Predicted RNA
pDC prec.
CITE-seq YS
LR probability predicted
1.0
.0

**Fig. S1.**

**Fig. S1: A single cell atlas of the human yolk sac**

**(A-C)** UMAP visualisation of cells shown in **Fig. 1B** coloured according to yolk sac (YS) scRNA-seq dataset source (**A**), tissue compartment (**B**), and refined annotation (**C**). HSPC= haematopoietic stem/progenitor cell, CMP= common myeloid progenitor, MEMP= megakaryocyte-erythroid-mast cell progenitor, LMPP= lymphoid-primed multipotent progenitor, Prog.= progenitor, MOP= monocyte progenitor, ILC= innate lymphoid cell, NK= natural killer cell, pDC= plasmacytoid DC, pre.= precursor, DC= dendritic cell, Mac.= macrophage, Eo Baso= eosinophil basophil, MK= megakaryocyte, AEC= arteriolar endothelial cell, HE= hemogenic endothelium, EC= endothelial cell (**Table S5**).

**(D)** FACS gating strategy used sort YS CD45$^{+/-}$ fractions for plate-based scRNA-seq. Representative gating from n=2 independent samples (5-7PCW).

(**E**) UMAP visualisation of the YS cells shown in **Fig. 1B** (n=10, k=169,798) integrated with k=143 cells from plate-based sequencing (SS2) cells (triangles) FACS isolated from n=2 individual donors (5-7PCW). Left: colours indicate cell states in droplet based scRNA-seq (10x). Right; colours indicate predicted cell states in plate-based scRNA-seq (SS2) (**Table S5, S8**).

(**F**) UMAP visualisation of cells sequenced using CITE-seq from n=2 biologically independent YS samples (k=3,559). Colours represent cell states (**Table S4, S9, S5**).

**(G)** Median logistic regression class assignment probabilities for a model trained on YS scRNA-seq cell states from **Fig. 1B** (y-axis) projected onto corresponding YS CITE-seq acquired cell states(x-axis) (**Table S31**).

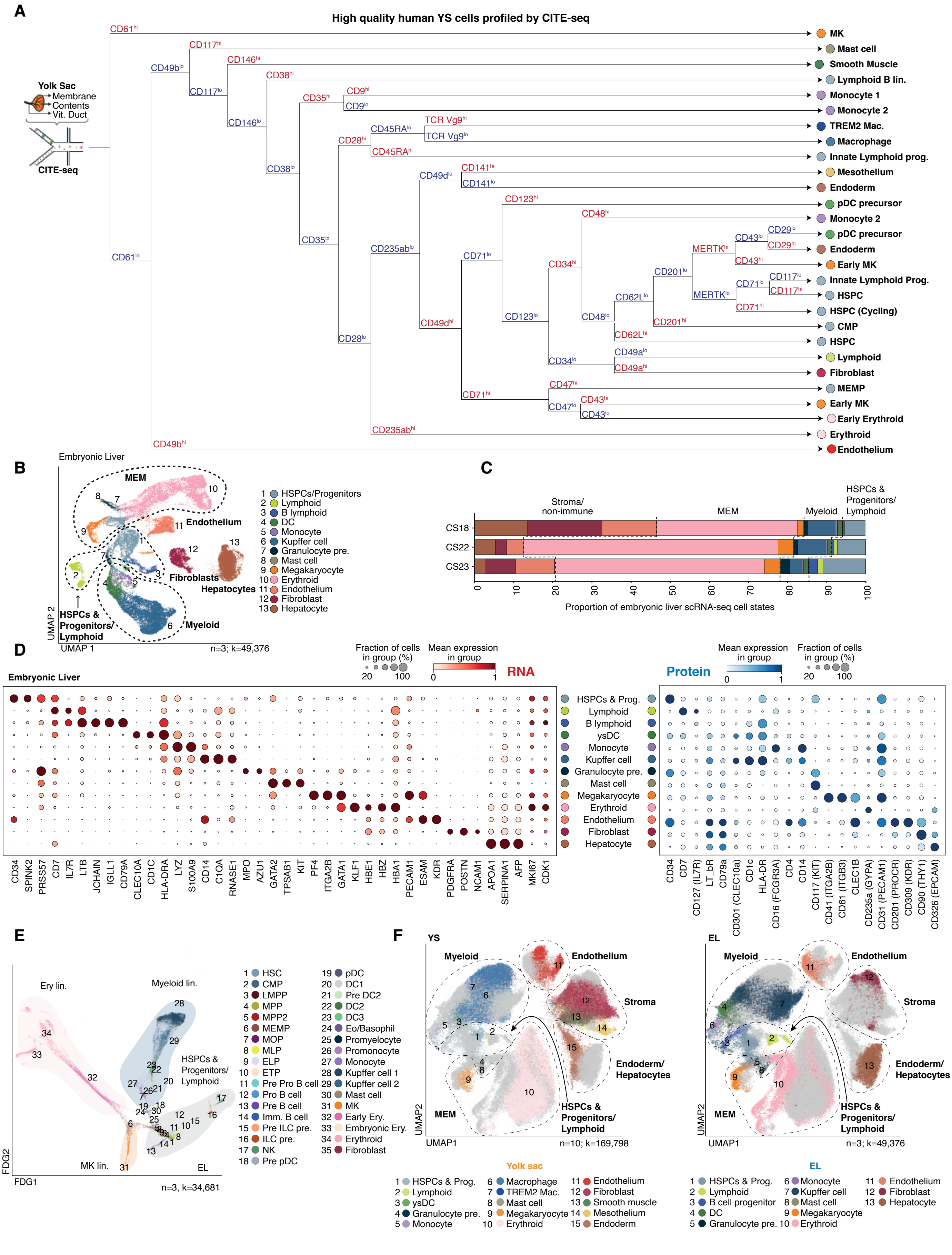
A
High quality human YS cells profiled by CITE-seq
Yolk Sac
Membrane
Contents
Vit. Duct
CITE-seq
CD61hi
CD61lo
CD49blo
CD49bhi
CD117hi
CD117lo
CD146hi
CD146lo
CD38hi
CD38lo
CD35hi
CD35lo
CD9hi
CD9lo
CD28hi
CD28lo
CD45RAlo
CD45RAhi
TCR Vg9hi
TCR Vg9lo
CD49dlo
CD49dhi
CD141hi
CD141lo
CD235ablo
CD235abhi
CD123hi
CD123lo
CD71lo
CD71hi
CD48hi
CD48lo
CD34hi
CD34lo
MERTKhi
MERTKlo
CD43lo
CD43hi
CD29lo
CD29hi
CD201lo
CD201hi
CD62Llo
CD62Lhi
CD117lo
CD117hi
CD49alo
CD49ahi
CD47hi
CD47lo
MK
Mast cell
Smooth Muscle
Lymphoid B lin.
Monocyte 1
Monocyte 2
TREM2 Mac.
Macrophage
Innate Lymphoid prog.
Mesothelium
Endoderm
pDC precursor
Monocyte 2
pDC precursor
Endoderm
Early MK
Innate Lymphoid Prog.
HSPC
HSPC (Cycling)
CMP
HSPC
Lymphoid
Fibroblast
MEMP
Early MK
Early Erythroid
Erythroid
Endothelium
B
Embryonic Liver
MEM
Endothelium
Fibroblasts
Hepatocytes
HSPCs & Progenitors/ Lymphoid
Myeloid
UMAP 2
UMAP 1
n=3; k=49,376
1 HSPCs/Progenitors
2 Lymphoid
3 B lymphoid
4 DC
5 Monocyte
6 Kupffer cell
7 Granulocyte pre.
8 Mast cell
9 Megakaryocyte
10 Erythroid
11 Endothelium
12 Fibroblast
13 Hepatocyte
C
Stroma/ non-immune
MEM
Myeloid
HSPCs & Progenitors/ Lymphoid
CS18
CS22
CS23
0 10 20 30 40 50 60 70 80 90 100
Proportion of embryonic liver scRNA-seq cell states
D
Embryonic Liver
Fraction of cells in group (%)
20 100
Mean expression in group
0 1
RNA
Protein
HSPCs & Prog.
Lymphoid
B lymphoid
ysDC
Monocyte
Kupffer cell
Granulocyte pre.
Mast cell
Megakaryocyte
Erythroid
Endothelium
Fibroblast
Hepatocyte
CD34 SPINK2 PRSS57 CD7 IL7R LTB JCHAIN IGLL1 CD79A CLEC10A CD1C HLA-DRA LYZ S100A9 CD14 C1QA RNASE1 MPO AZU1 GATA2 TPSAB1 KIT PF4 ITGA2B GATA1 KLF1 HBE1 HBZ HBA1 PECAM1 ESAM KDR PDGFRA POSTN NCAM1 APOA1 SERPINA1 AFP MKI67 CDK1
CD34 CD7 CD127 (IL7R) LT_bR CD79a CD301 (CLEC10a) CD1c HLA-DR CD16 (FCGR3A) CD4 CD14 CD117 (KIT) CD41 (ITGA2B) CD61 (ITGB3) CLEC1B CD235a (GYPA) CD31 (PECAM1) CD201 (PROCR) CD309 (KDR) CD90 (THY1) CD326 (EPCAM)
E
Ery lin.
Myeloid lin.
HSPCs & Progenitors/ Lymphoid
MK lin.
EL
FDG2
FDG1
n=3, k=34,681
1 HSC
2 CMP
3 LMPP
4 MPP
5 MPP2
6 MEMP
7 MOP
8 MLP
9 ELP
10 ETP
11 Pre Pro B cell
12 Pro B cell
13 Pre B cell
14 Imm. B cell
15 Pre ILC pre.
16 ILC pre.
17 NK
18 Pre pDC
19 pDC
20 DC1
21 Pre DC2
22 DC2
23 DC3
24 Eo/Basophil
25 Promyelocyte
26 Promonocyte
27 Monocyte
28 Kupffer cell 1
29 Kupffer cell 2
30 Mast cell
31 MK
32 Early Ery.
33 Embryonic Ery.
34 Erythroid
35 Fibroblast
F
YS
Myeloid
Endothelium
Stroma
Endoderm/ Hepatocytes
HSPCs & Progenitors/ Lymphoid
MEM
UMAP2
UMAP1
n=10; k=169,798
EL
Myeloid
Endothelium
Stroma
Endoderm/ Hepatocytes
HSPCs & Progenitors/ Lymphoid
MEM
UMAP2
UMAP1
n=3; k=49,376
Yolk sac
1 HSPCs & Prog.
2 Lymphoid
3 ysDC
4 Granulocyte pre.
5 Monocyte
6 Macrophage
7 TREM2 Mac.
8 Mast cell
9 Megakaryocyte
10 Erythroid
11 Endothelium
12 Fibroblast
13 Smooth muscle
14 Mesothelium
15 Endoderm
EL
1 HSPCs & Prog.
2 Lymphoid
3 B cell progenitor
4 DC
5 Granulocyte pre.
6 Monocyte
7 Kupffer cell
8 Mast cell
9 Megakaryocyte
10 Erythroid
11 Endothelium
12 Fibroblast
13 Hepatocyte

**Fig. S2.**

**Fig. S2: A single cell atlas of the human yolk sac**

**(A)** Decision tree showing the step-wise relative importance (left to right) and expression (hi/lo) of protein markers in determining YS cell states using the protein data from YS CITE-seq. The optimum number of splits was determined by minimum combination of error rate and standard deviation scores whilst n_splits being > number of cell states ingested, resulting in 27 branches (**Table S4**).

**(B)** UMAP visualisation of the cell states identified in the embryonic liver (EL) scRNA-seq dataset from n=3 independent biological repeats (k=49,376). Colours represent cell states. DC= dendritic cell, MEM= megakaryocyte-erythroid-mast cell lineage, pre.= precursor.

**(C)** Stacked bar plot displaying the proportion of each cell state found in each embryonic liver scRNA-seq data sample. Colours match cell states in **B.**

**(D)** Dot plot showing the expression level (by colour) and percent expression (by dot size) of broad cell state-defining genes in scRNA-seq data (n=3, k=49,376) (left, log-normalised and standard_scale='var'), and their protein counterparts in the liver CITE-seq dataset (n=9, k=57,310) (right, data scaled zero_centre = False, standard_scale='var') (**Table S4**).

**(E)** Force directed graph (FDG) visualisation of the haematopoietic cell states identified in the EL scRNA-seq from n=3 biologically independent donors (k=34,681). Colours represent cell states and clouds represent lineages. CMP= common myeloid progenitor, DC= dendritic cell, ELP= early lymphoid progenitor, Eo/Baso= eosinophil/basophil, Ery= erythroid, ETP= early thymic progenitor, HE= hemogenic endothelium, HSC= haematopoietic stem cell, HSPC= haematopoietic stem progenitor cell; ILC, innate lymphoid cell; LMPP, lymphoid-primed multipotent progenitor; Mac, macrophage; MEM, megakaryocyte-erythroid-mast cell lineage; MEMP, megakaryocyte-erythroid-mast cell progenitor; MK, megakaryocyte; MLP, multi-lymphoid progenitor; Mono, monocyte; MOP, monocyte progenitor; MPP, multipotent progenitor; Neut, neutrophil; NK, natural killer cell; pDC, plasmacytoid DC; pre., precursor; prog., progenitor; prolif., proliferating (**Table S5**).

**(F)** UMAP visualisation of the merged YS and EL scRNA-seq data shown in **Fig. 1B** and **Fig. S3B** respectively and coloured by cell state and tissue (Left, YS, n=10, k=169,798; Right EL, n=3, k=49,376) (**Table S5**).

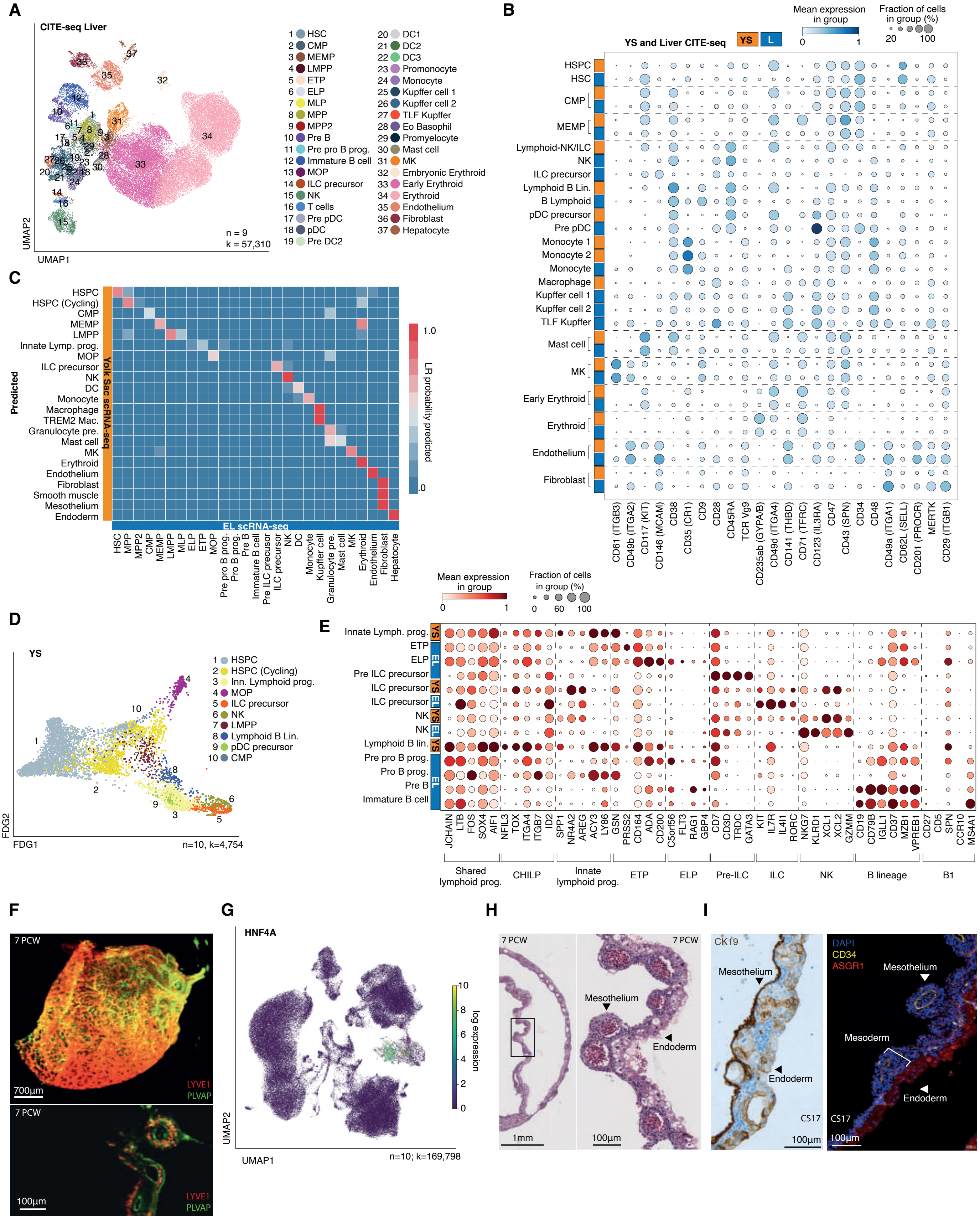
A
CITE-seq Liver
UMAP2
UMAP1
n = 9
k = 57,310
1 HSC
2 CMP
3 MEMP
4 LMPP
5 ETP
6 ELP
7 MLP
8 MPP
9 MPP2
10 Pre B
11 Pre pro B prog.
12 Immature B cell
13 MOP
14 ILC precursor
15 NK
16 T cells
17 Pre pDC
18 pDC
19 Pre DC2
20 DC1
21 DC2
22 DC3
23 Promonocyte
24 Monocyte
25 Kupffer cell 1
26 Kupffer cell 2
27 TLF Kupffer
28 Eo Basophil
29 Promyelocyte
30 Mast cell
31 MK
32 Embryonic Erythroid
33 Early Erythroid
34 Erythroid
35 Endothelium
36 Fibroblast
37 Hepatocyte
B
YS and Liver CITE-seq
YS
L
Mean expression in group
Fraction of cells in group (%)
HSPC
HSC
CMP
MEMP
Lymphoid-NK/ILC
NK
ILC precursor
Lymphoid B Lin.
B Lymphoid
pDC precursor
Pre pDC
Monocyte 1
Monocyte 2
Monocyte
Macrophage
Kupffer cell 1
Kupffer cell 2
TLF Kupffer
Mast cell
MK
Early Erythroid
Erythroid
Endothelium
Fibroblast
CD61 (ITGB3)
CD49b (ITGA2)
CD117 (KIT)
CD146 (MCAM)
CD38
CD35 (CR1)
CD9
CD28
CD45RA
TCR Vg9
CD235ab (GYPA/B)
CD49d (ITGA4)
CD141 (THBD)
CD71 (TFRC)
CD123 (IL3RA)
CD47
CD43 (SPN)
CD34
CD48
CD49a (ITGA1)
CD62L (SELL)
CD201 (PROCR)
MERTK
CD29 (ITGB1)
C
Predicted
Yolk Sac scRNA-seq
EL scRNA-seq
LR probability predicted
HSPC
HSPC (Cycling)
CMP
MEMP
LMPP
Innate Lymp. prog.
MOP
ILC precursor
NK
DC
Monocyte
Macrophage
TREM2 Mac.
Granulocyte pre.
Mast cell
MK
Erythroid
Endothelium
Fibroblast
Smooth muscle
Mesothelium
Endoderm
HSC
MPP
MPP2
CMP
MEMP
LMPP
MLP
ELP
ETP
MOP
Pre pro B prog.
Pro B prog.
Pre B
Immature B cell
Pre ILC precursor
ILC precursor
NK
DC
Monocyte
Kupffer cell
Granulocyte pre.
Mast cell
MK
Erythroid
Endothelium
Fibroblast
Hepatocyte
D
YS
FDG2
FDG1
n=10, k=4,754
1 HSPC
2 HSPC (Cycling)
3 Inn. Lymphoid prog.
4 MOP
5 ILC precursor
6 NK
7 LMPP
8 Lymphoid B Lin.
9 pDC precursor
10 CMP
E
Mean expression in group
Fraction of cells in group (%)
Innate Lymph. prog.
ETP
ELP
Pre ILC precursor
ILC precursor
ILC precursor
NK
NK
Lymphoid B lin.
Pre pro B prog.
Pro B prog.
Pre B
Immature B cell
JCHAIN
LTB
FOS
SOX4
AIF1
NFIL3
TOX
ITGA4
ITGB7
ID2
SPP1
NR4A2
AREG
ACY3
LY86
GSN
PRSS2
CD164
ADA
CD200
C5orf56
FLT3
RAG1
GBP4
CD7
CD3D
TRDC
GATA3
KIT
IL7R
IL4I1
RORC
NKG7
KLRD1
XCL1
XCL2
GZMM
CD19
CD79B
IGLL1
CD37
MZB1
VPREB1
CD27
CD5
SPN
CCR10
MS4A1
Shared lymphoid prog.
CHILP
Innate lymphoid prog.
ETP
ELP
Pre-ILC
ILC
NK
B lineage
B1
F
7 PCW
700µm
LYVE1
PLVAP
7 PCW
100µm
LYVE1
PLVAP
G
HNF4A
log expression
UMAP2
UMAP1
n=10; k=169,798
H
7 PCW
1mm
7 PCW
Mesothelium
Endoderm
100µm
I
CK19
Mesothelium
Endoderm
CS17
100µm
DAPI
CD34
ASGR1
Mesothelium
Mesoderm
Endoderm
CS17
100µm

**Fig. S3.**

**Fig. S3: A single cell atlas of the human yolk sac**

**(A)** UMAP visualisation of cells sequenced using liver CITE-seq from n=9 biologically independent liver samples (k=57,310). Coloured by individual cell state. HSPC= haematopoietic stem/progenitor cell, CMP= common myeloid progenitor, MEMP= megakaryocyte-erythroid-mast cell progenitor, LMPP= lymphoid-primed multipotent progenitor,ETP= Early Thymic progenitor, ELP= Early Lymphoid progenitor, MLP= Multi-potent Lymphoid progenitor, MPP/MPP2= Multipotent progenitor (2), Prog.= progenitor, MOP= monocyte progenitor, ILC= innate lymphoid cell, NK= natural killer cell, pDC= plasmacytoid DC, pre.= precursor, DC= dendritic cell, Mac.= macrophage, Eo Baso= eosinophil basophil, MK= megakaryocyte, AEC= arteriolar endothelial cell, HE= hemogenic endothelium, EC= endothelial cell (**Table S30, S5**).

**(B)** Dot plot showing the level (colour scale) and percent expression (dot size) of select proteins per matched refined cell state (including those shown in the decision tree in (**A**) in the YS and liver CITE-seq datasets. HSC, haematopoietic stem cell; HSPC, haematopoietic stem/ progenitor cell; CMP, common myeloid progenitor; MEMP, megakaryocyte-erythroid-mast cell progenitor; pDC, plasmacytoid dendritic cell; MK, megakaryocyte (data scaled zero_centre = False) (**Table S9, S30**).

**(C)** Median logistic regression class prediction probabilities for a model trained on embryonic liver (EL) scRNA-seq cell states (x-axis) and projected onto corresponding YS scRNA-seq acquired cell states (y-axis) (**Table S12**).

(**D**) Force directed graph (FDG) visualisation of lymphoid cell states in the YS scRNA-seq dataset (n=10, k=4,754) (**Table S5**).

**(E)** Dot plot showing the expression level (colour scale) and percent expression (dot size) of lymphoid population-defining gene markers in lymphoid lineage cell states in the YS and EL scRNA-seq datasets.

**(F)** Stills of 3D (top) and 2D z-stack (bottom) light-sheet fluorescence microscopy images of 7PCW YS stained with LYVE1 (red) and PVLAP (green). Inset, a zoom in showing sinusoidal architecture. Top scale bar=700µm, bottom scale bar=100µm; see **Movie S2-S3.**

**(G)** Feature plot of YS scRNA-seq data (as shown in **Fig. 1B**) log-normalised and scaled max value=10 expression of *HNF4A* (n=10, k=169,798) (**Table S5**).

**(H)** H&E staining of a 7PCW YS, representative image from one of n=4 biologically independent samples (4-8PCW), with mesothelium and endoderm marked by arrows. Scale bar=1mm (left), 100µm (right).

**(I)** Left: CK19 staining of a 7PCW YS, representative image from one of n=4 biologically independent samples. Scale bar=100µm. Right: Immunofluorescence staining of a 7PCW YS for endoderm with ASGR1 (red), endothelium with CD34 (yellow) and DAPI-rich (blue) mesoderm. Scale bar=100µm.

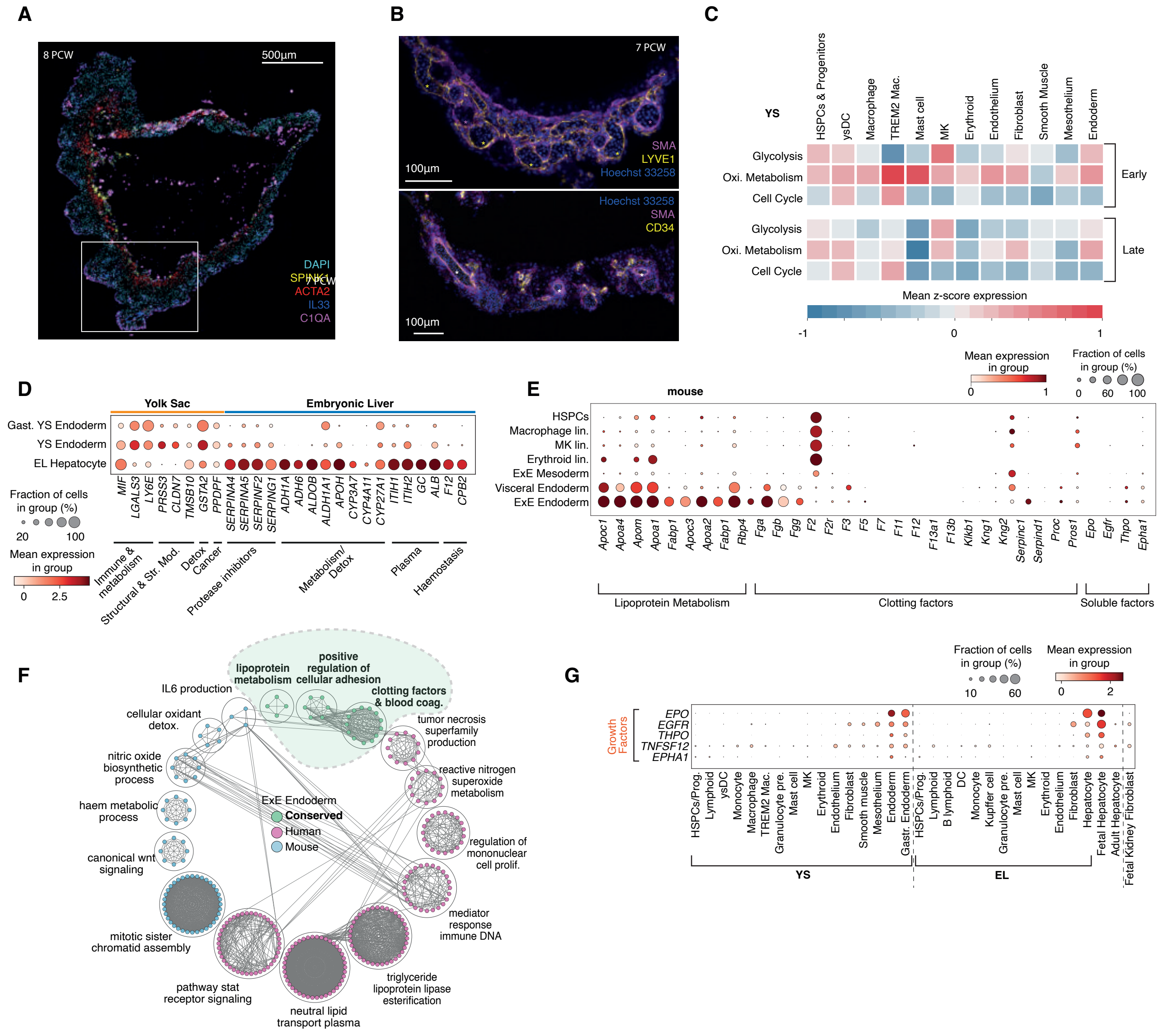

A
8 PCW
500μm
7 PCW
DAPI
SPINK1
ACTA2
IL33
C1QA
B
7 PCW
SMA
LYVE1
Hoechst 33258
100μm
Hoechst 33258
SMA
CD34
100μm
C
YS
HSPCs & Progenitors
ysDC
Macrophage
TREM2 Mac.
Mast cell
MK
Erythroid
Endothelium
Fibroblast
Smooth Muscle
Mesothelium
Endoderm
Glycolysis
Oxi. Metabolism
Cell Cycle
Early
Late
Mean z-score expression
-1
0
1
D
Yolk Sac
Embryonic Liver
Gast. YS Endoderm
YS Endoderm
EL Hepatocyte
Fraction of cells in group (%)
20
100
Mean expression in group
0
2.5
MIF LGALS3 LY6E PRSS3 CLDN7 TMSB10 GSTA2 PPDPF SERPINA4 SERPINA5 SERPINF2 SERPING1 ADH1A ADH6 ALDOB ALDH1A1 APOH CYP3A7 CYP4A11 CYP27A1 ITIH1 ITIH2 GC ALB F12 CPB2
Immune & metabolism
Structural & Str. Mod.
Detox
Cancer
Protease inhibitors
Metabolism/ Detox
Plasma
Haemostasis
E
mouse
Mean expression in group
0
1
Fraction of cells in group (%)
0
60
100
HSPCs
Macrophage lin.
MK lin.
Erythroid lin.
ExE Mesoderm
Visceral Endoderm
ExE Endoderm
Apoc1 Apoa4 Apom Apoa1 Fabp1 Apoc3 Apoa2 Fabp1 Rbp4 Fga Fgb Fgg F2 F2r F3 F5 F7 F11 F12 F13a1 F13b Klkb1 Kng1 Kng2 Serpinc1 Serpind1 Proc Pros1 Epo Egfr Thpo Epha1
Lipoprotein Metabolism
Clotting factors
Soluble factors
F
positive regulation of cellular adhesion
lipoprotein metabolism
clotting factors & blood coag.
IL6 production
cellular oxidant detox.
tumor necrosis superfamily production
nitric oxide biosynthetic process
reactive nitrogen superoxide metabolism
haem metabolic process
ExE Endoderm
Conserved
Human
Mouse
regulation of mononuclear cell prolif.
canonical wnt signaling
mediator response immune DNA
mitotic sister chromatid assembly
pathway stat receptor signaling
triglyceride lipoprotein lipase esterification
neutral lipid transport plasma
G
Fraction of cells in group (%)
10
60
Mean expression in group
0
2
Growth Factors
EPO
EGFR
THPO
TNFSF12
EPHA1
HSPCs/Prog. Lymphoid ysDC Monocyte Macrophage TREM2 Mac. Granulocyte pre. Mast cell MK Erythroid Endothelium Fibroblast Smooth muscle Mesothelium Endoderm Gastr. Endoderm
HSPCs/Prog. Lymphoid B lymphoid DC Monocyte Kupffer cell Granulocyte pre. Mast cell MK Erythroid Endothelium Fibroblast Hepatocyte
Fetal Hepatocyte
Adult Hepatocyte
Fetal Kidney Fibroblast
YS
EL

**Fig. S4.**

**Fig. S4: A single cell atlas of the human yolk sac & Multiorgan functions of YS**

**(A)** RNAscope imaging of an 8PCW YS for Endoderm (*SPINK1*), Smooth muscle (*ACTA2*), AEC (*IL33*), Macrophage (*C1QA*) with DAPI. White box indicates ROI shown in **Fig. 1E**. Scale bar=500um.

**(B)** A 7PCW YS stained with Hoechst 33268 (blue), SMA (magenta) and CD34 (yellow; top) or LYVE1 (yellow, bottom) and imaged using confocal microscopy. Scale bars=100μm.

**(C)** Heatmap of z-normalised GO geneset module scores between early and late predicted Milo neighbourhoods for cell cycle (GO:0022402), oxidative metabolism (GO:0045333), and glycolysis (GO:0006096), subtracted by the mean expression of 50 randomly sampled genes at 25 bins using *scanpy.tl.score_genes*.

**(D)** Dot plot showing the expression level (colour scale) and percent expression (dot size) of selected DEGs (**Table S3, S20, S7**) between YS endoderm (main and gastrula (gast.) data) and embryonic liver (EL) hepatocytes (data scaled max_value=10, gastrulation data scaled independently).

**(E)** Dot plot showing the expression level (colour scale) and percent expression (dot size) of clotting and soluble factors in selected mouse gastrulation (*56*) cell states.

**(F)** Flower plot of the significant genesets enriched in YS endoderm (pink), mouse extraembryonic (ExE) endoderm (blue) and conserved between species (green). Nodes indicate significantly enriched gene sets (Q-value <0.05) whilst edges between nodes represent gene overlap between gene sets. Annotated grouping circles indicate markov cluster neighbourhoods of gene expression modules which share high gene set similarities (**Table S21**).

**(G)** Dot plot showing the expression level (colour scale) and percent expression (dot size) of soluble factors in YS (main and gastrula), EL, fetal/adult liver and fetal kidney scRNA-seq select cell states (each dataset scaled max_value=10 independently then combined except YS and EL scRNA-seq).

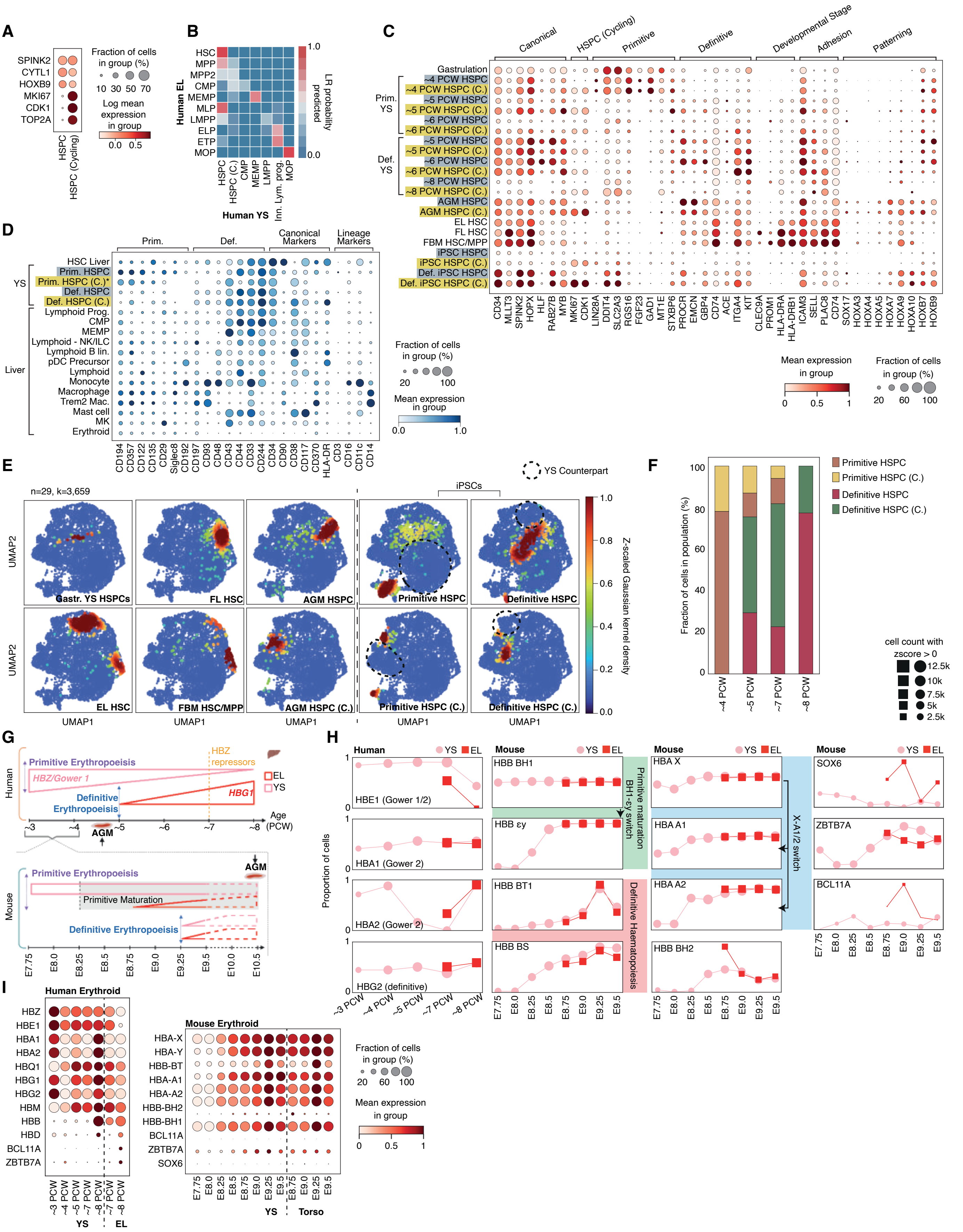
A
SPINK2
CYTL1
HOXB9
MKI67
CDK1
TOP2A
HSPC
HSPC (Cycling)
Fraction of cells in group (%)
10 30 50 70
Log mean expression in group
0.0 0.5
B
Human EL
HSC
MPP
MPP2
CMP
MEMP
MLP
LMPP
ELP
ETP
MOP
HSPC
HSPC (C.)
CMP
MEMP
LMPP
Inn. Lym. prog.
MOP
Human YS
LR probability predicted
1.0
0.0
C
Canonical
HSPC (Cycling)
Primitive
Definitive
Developmental Stage
Adhesion
Patterning
Gastrulation
~4 PCW HSPC
~4 PCW HSPC (C.)
~5 PCW HSPC
~5 PCW HSPC (C.)
~6 PCW HSPC
~6 PCW HSPC (C.)
~5 PCW HSPC
~5 PCW HSPC (C.)
~6 PCW HSPC
~6 PCW HSPC (C.)
~8 PCW HSPC
~8 PCW HSPC (C.)
AGM HSPC
AGM HSPC (C.)
EL HSC
FL HSC
FBM HSC/MPP
iPSC HSPC
iPSC HSPC (C.)
Def. iPSC HSPC
Def. iPSC HSPC (C.)
Prim. YS
Def. YS
CD34 MLLT3 SPINK2 HOPX HLF RAB27B MYB MKI67 CDK1 LIN28A DDIT4 SLC2A3 RGS16 FGF23 GAD1 MT1E STXBP6 PROCR EMCN GBP4 CD74 ACE ITGA4 KIT CLEC9A PROM1 HLA-DRA HLA-DRB1 ICAM3 SELL PLAC8 CD74 SOX17 HOXA3 HOXA4 HOXA5 HOXA7 HOXA9 HOXA10 HOXB7 HOXB9
Mean expression in group
0 0.5 1
Fraction of cells in group (%)
20 60 100
D
Prim.
Def.
Canonical Markers
Lineage Markers
YS
HSC Liver
Prim. HSPC
Prim. HSPC (C.)*
Def. HSPC
Def. HSPC (C.)
Liver
Lymphoid Prog.
CMP
MEMP
Lymphoid - NK/ILC
Lymphoid B lin.
pDC Precursor
Lymphoid
Monocyte
Macrophage
Trem2 Mac.
Mast cell
MK
Erythroid
CD194 CD357 CD122 CD135 CD29 Siglec8 CD192 CD197 CD93 CD48 CD43 CD44 CD33 CD244 CD34 CD90 CD38 CD117 CD370 HLA-DR CD3 CD16 CD11c CD14
Fraction of cells in group (%)
20 100
Mean expression in group
0.0 1.0
E
n=29, k=3,659
YS Counterpart
iPSCs
UMAP2
UMAP1
Gastr. YS HSPCs
FL HSC
AGM HSPC
Primitive HSPC
Definitive HSPC
EL HSC
FBM HSC/MPP
AGM HSPC (C.)
Primitive HSPC (C.)
Definitive HSPC (C.)
Z-scaled Gaussian kernel density
1.0 0.8 0.6 0.4 0.2 0.0
F
Primitive HSPC
Primitive HSPC (C.)
Definitive HSPC
Definitive HSPC (C.)
Fraction of cells in population (%)
100 80 60 40 20 0
~4 PCW
~5 PCW
~7 PCW
~8 PCW
cell count with zscore > 0
12.5k
10k
7.5k
5k
2.5k
G
Human
Primitive Erythropoeisis
HBZ/Gower 1
HBZ repressors
EL
YS
Definitive Erythropoeisis
HBG1
AGM
~3 ~4 ~5 ~6 ~7 ~8
Age (PCW)
Mouse
Primitive Erythropoeisis
AGM
Primitive Maturation
Definitive Erythropoeisis
E7.75 E8.0 E8.25 E8.5 E8.75 E9.0 E9.25 E9.5 E10.0 E10.5
H
Human
YS
EL
Mouse
Proportion of cells
HBE1 (Gower 1/2)
HBA1 (Gower 2)
HBA2 (Gower 2)
HBG2 (definitive)
~3 PCW ~4 PCW ~5 PCW ~7 PCW ~8 PCW
HBB BH1
HBB εy
HBB BT1
HBB BS
Primitive maturation BH1-εy switch
Definitive Haematopoiesis
HBA X
HBA A1
HBA A2
HBB BH2
X-A1/2 switch
SOX6
ZBTB7A
BCL11A
E7.75 E8.0 E8.25 E8.5 E8.75 E9.0 E9.25 E9.5
I
Human Erythroid
HBZ
HBE1
HBA1
HBA2
HBQ1
HBG1
HBG2
HBM
HBB
HBD
BCL11A
ZBTB7A
~3 PCW ~4 PCW ~5 PCW ~7 PCW ~8 PCW ~7 PCW ~8 PCW
YS
EL
Mouse Erythroid
HBA-X
HBA-Y
HBB-BT
HBA-A1
HBA-A2
HBB-BH2
HBB-BH1
BCL11A
ZBTB7A
SOX6
E7.75 E8.0 E8.25 E8.5 E8.75 E9.0 E9.25 E9.5 E8.75 E9.0 E9.25 E9.5
YS
Torso
Fraction of cells in group (%)
20 60 100
Mean expression in group
0 0.5 1

**Fig. S5.**

**Fig. S5: Primitive versus definitive haematopoiesis in YS and liver**

**(A)** Dot plot showing the expression level (colour scale) and percent expression (dot size) of genes distinguishing YS HSPC from YS cycling HSPC in the YS (main) scRNA-seq data (data scaled max_value=10).

**(B)** Median logistic regression class prediction probabilities for a model trained on EL progenitor scRNA-seq cell states (x-axis) projected onto YS scRNA-seq cell states (y-axis)(**Table S12**).

**(C)** Dot plot showing the mean variance scaled expression (by colour) and percentage of population expressing (by dot size) of canonical, cycling HSPC-specific, primitive, definitive, developmental-stage specific, adhesion and patterning HSC markers expressed between YS HSPCs (split by HSPC/ cycling HSPC and primitive/definitive) across time including gastrulation (67), AGM HSPC (66), matched EL HSC, FL HSC (*9*), fetal BM HSC/MPP (*54*), iPSC-derived HSPC (*21*) and definitive iPSC-derived HPSC (*11*)**.**

**(D)** Dot plot showing the expression level (by colour) and percent expression (by dot size) of differentially expressed proteins between primitive and definitive HSPCS, alongside canonical HSC markers and lineage markers, in selected cell states from YS and liver CITE-seq data (datasets scaled zero_centre=False independently then standard_scale='var').

**(E)** Density plots showing the distribution of indicated HSPC population in the integrated UMAP landscape of HSPC/HSCs from YS (n=10, k=2,597), YS gastrulation (*57*) (n=1, k=23), AGM (*58*) (n=3, k=182), matched embryonic liver (EL) (n=3, k=412), fetal liver (FL) (n=14, k=242), fetal bone marrow (*54*) (FBM) (n=9, k=92), iPSC-derived HSPC (n=12, k=355) (*21*) and definitive iPSC-derived HSPC (n=2, k=273) (*11*). scRNA-seq datasets. The colour of HSC/HSPC cells represents the z-scored kernel density estimation (KDE) score for each population (**Table S5**).

**(F)** Bar graph showing the proportional representation of primitive YS HSPC and cycling HSPC to definitive YS HSPC and cycling HSPC in the main and gastrulation YS scRNA-seq data (grouped by gestational age in PCW).

**(G)** Schematic showing the relative timescales of primitive and definitive erythropoiesis in human and mouse, and contributions of AGM, EL, and YS to this process.

**(H)** Left: Line graph showing the relative change in expression of Gower 1/2 globin *HBE1*, Gower 2 globins *HBA1/2* and definitive globin *HBG2* in human erythroid cells from YS (pink) and matched embryonic liver (EL; red) over gestational age. Middle and right: Globin expression in mouse erythroid cells (*56*) including HBB BH1, εγ, X, A2, BT1 and BS. Pink lines= mouse YS scRNA-seq data. Red lines= aged-matched mouse torso scRNA-seq data. Mouse haemoglobins implicated in primitive maturation, definitive haematopoiesis, and a switch between the two are grouped. The y-axis represents the proportion of erythroid lineage cells.

**(I)** Dot plot showing the standard scaled (max value=1) expression level (colour scale) and percent expression (dot size) of Hb genes expressed in human YS scRNA-seq (main and gastrulation) and mouse YS scRNA-seq data (*56*) erythroid lineage cells by gestational age.

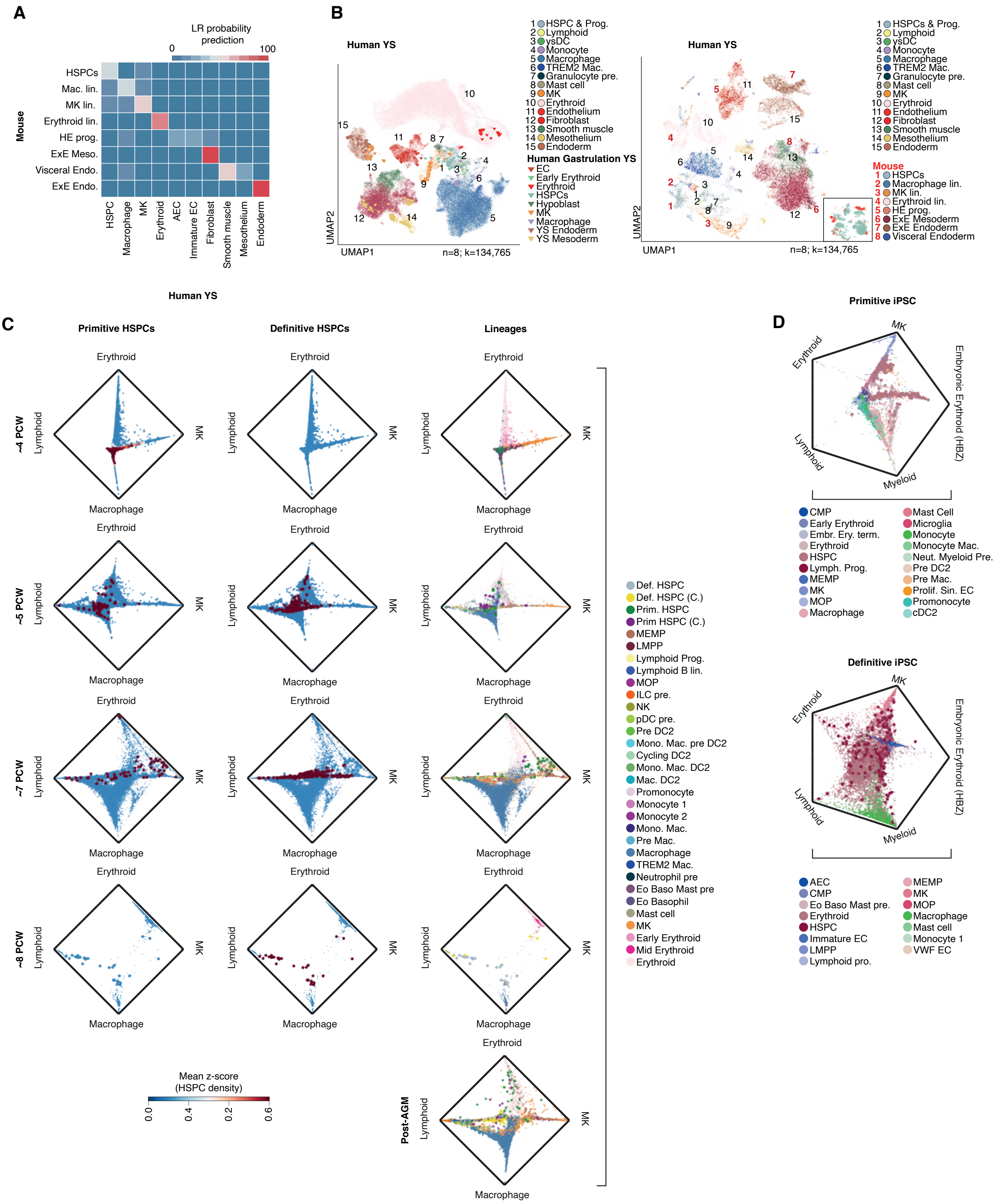
A
LR probability prediction
0
100
Mouse
HSPCs
Mac. lin.
MK lin.
Erythroid lin.
HE prog.
ExE Meso.
Visceral Endo.
ExE Endo.
HSPC
Macrophage
MK
Erythroid
AEC
Immature EC
Fibroblast
Smooth muscle
Mesothelium
Endoderm
Human YS
B
Human YS
UMAP2
UMAP1
n=8; k=134,765
1 HSPC & Prog.
2 Lymphoid
3 ysDC
4 Monocyte
5 Macrophage
6 TREM2 Mac.
7 Granulocyte pre.
8 Mast cell
9 MK
10 Erythroid
11 Endothelium
12 Fibroblast
13 Smooth muscle
14 Mesothelium
15 Endoderm
Human Gastrulation YS
EC
Early Erythroid
Erythroid
HSPCs
Hypoblast
MK
Macrophage
YS Endoderm
YS Mesoderm
Human YS
UMAP2
UMAP1
n=8; k=134,765
1 HSPCs & Prog.
2 Lymphoid
3 ysDC
4 Monocyte
5 Macrophage
6 TREM2 Mac.
7 Granulocyte pre.
8 Mast cell
9 MK
10 Erythroid
11 Endothelium
12 Fibroblast
13 Smooth muscle
14 Mesothelium
15 Endoderm
Mouse
1 HSPCs
2 Macrophage lin.
3 MK lin.
4 Erythroid lin.
5 HE prog.
6 ExE Mesoderm
7 ExE Endoderm
8 Visceral Endoderm
C
Primitive HSPCs
Definitive HSPCs
Lineages
~4 PCW
~5 PCW
~7 PCW
~8 PCW
Post-AGM
Erythroid
Lymphoid
MK
Macrophage
Mean z-score (HSPC density)
0.0
0.4
0.2
0.6
Def. HSPC
Def. HSPC (C.)
Prim. HSPC
Prim HSPC (C.)
MEMP
LMPP
Lymphoid Prog.
Lymphoid B lin.
MOP
ILC pre.
NK
pDC pre.
Pre DC2
Mono. Mac. pre DC2
Cycling DC2
Mono. Mac. DC2
Mac. DC2
Promonocyte
Monocyte 1
Monocyte 2
Mono. Mac.
Pre Mac.
Macrophage
TREM2 Mac.
Neutrophil pre
Eo Baso Mast pre
Eo Basophil
Mast cell
MK
Early Erythroid
Mid Erythroid
Erythroid
Primitive iPSC
D
Erythroid
MK
Embryonic Erythroid (HBZ)
Lymphoid
Myeloid
CMP
Early Erythroid
Embr. Ery. term.
Erythroid
HSPC
Lymph. Prog.
MEMP
MK
MOP
Macrophage
Mast Cell
Microglia
Monocyte
Monocyte Mac.
Neut. Myeloid Pre.
Pre DC2
Pre Mac.
Prolif. Sin. EC
Promonocyte
cDC2
Definitive iPSC
Erythroid
MK
Embryonic Erythroid (HBZ)
Lymphoid
Myeloid
AEC
CMP
Eo Baso Mast pre.
Erythroid
HSPC
Immature EC
LMPP
Lymphoid pro.
MEMP
MK
MOP
Macrophage
Mast cell
Monocyte 1
VWF EC

**Fig. S6.**

**Fig. S6: Primitive versus definitive haematopoiesis in YS and liver**

**(A)** Median logistic regression class prediction probabilities for a model trained on human YS scRNA-seq cell states (x-axis) projected onto corresponding mouse extraembryonic cell states from mouse gastrulation dataset (*56*) (y-axis). Prog.= progenitor, AEC= arteriolar endothelial cell, EC= endothelial cell, HSPC= haematopoietic stem and progenitor cell (**Table S13**).

**(B)** Left: UMAP visualisation of matched haematopoietic cell states in human YS scRNA-seq (dots) as shown in **Fig. 1B** (n=10, k=169,798) integrated with human gastrulation (CS7) scRNA-seq data (*57*) (triangles) (n=1, k=91). Lin.= lineage, pre.= precursor, DC= dendritic cell, MK= megakaryocyte, EC= endothelial cell (**Table S5)**. Right: UMAP visualisation of human YS scRNA-seq (as shown in **Fig. 1B**) (n=10, k=169,798) and equivalent mouse gastrulation extraembryonic cell states (*56*) (n=36, k=139,331). Insert highlighting location of mouse cell states within UMAP. Colours represent cell states. ExE,= extra embryonic, lin.= lineage, HE= hemogenic endothelium, and inset coloured by species (mouse= red, human= teal).

**(C)** Circular plots showing relative absorption probabilities of lineage-state transition between primitive (left), definitive HSPCs (middle) and lineage-specific cell state annotations (right) across YS gestational stages between CS10-11, CS14-15, CS17-18 and CS22-23. Colour indicates the HSPC population density as a z-scored kernel density estimation (KDE) score and the position of HSPC population densities indicate respective lineage priming probability between Macrophage, lymphoid (NK and B lineage), erythroid and MK terminal states.

**(D)** Circular plots showing relative probabilities of lineage-state transition between iPSC-derived HSPCs from the primitive culture protocol (top) and iPSC-derived HSPCs from the definitive culture protocol (bottom). Colour indicates the cell state annotation and the position of each population indicates respective lineage priming probability between myeloid, lymphoid erythroid, embryonic erythroid and MK terminal states.

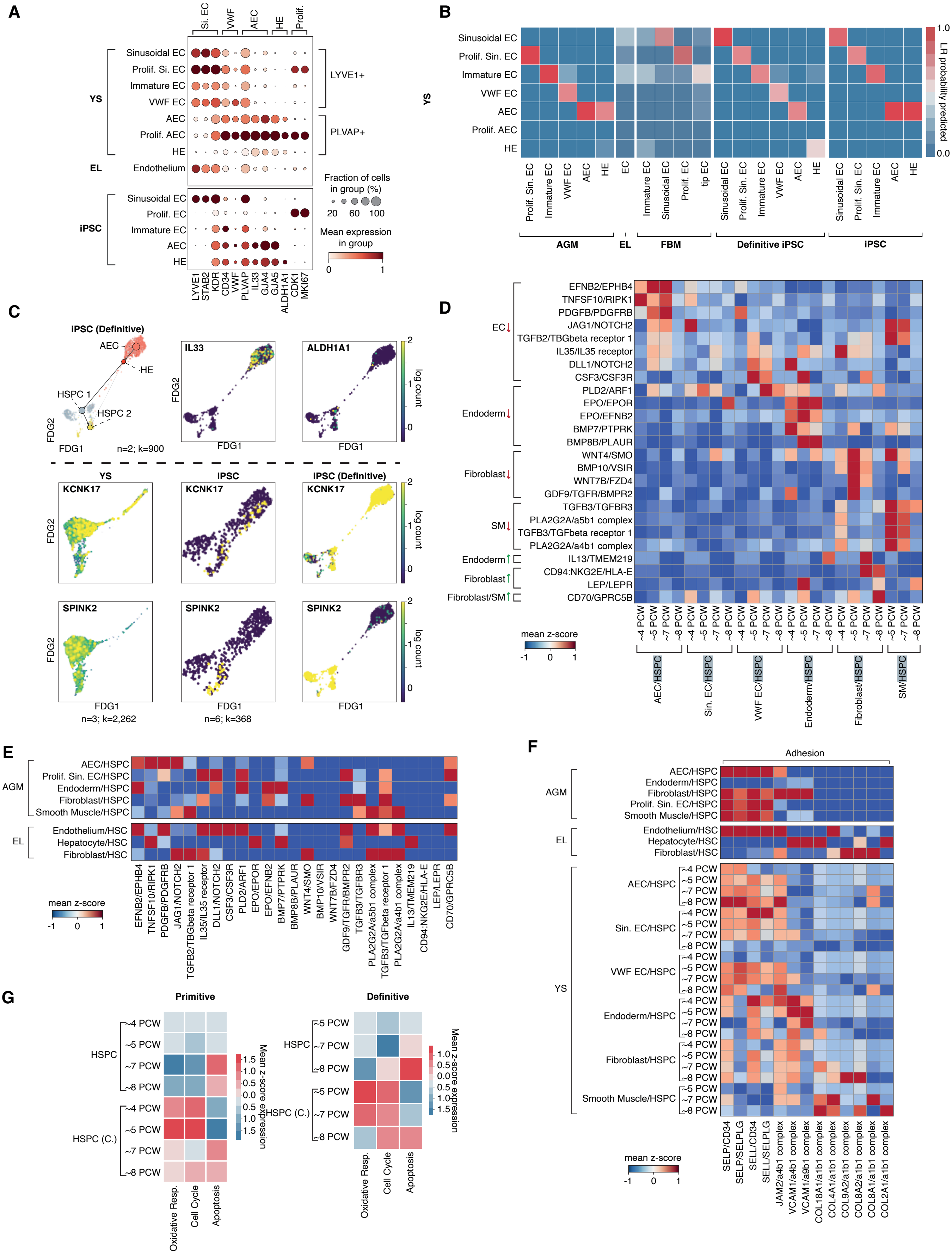
A
Si. EC
VWF
AEC
HE
Prolif.
Sinusoidal EC
Prolif. Si. EC
Immature EC
VWF EC
AEC
Prolif. AEC
HE
Endothelium
YS
EL
LYVE1+
PLVAP+
Fraction of cells in group (%)
20 60 100
Mean expression in group
0 1
iPSC
Sinusoidal EC
Prolif. EC
Immature EC
AEC
HE
LYVE1 STAB2 KDR CD34 VWF PLVAP IL33 GJA4 GJA5 ALDH1A1 CDK1 MKI67
B
Sinusoidal EC
Prolif. Sin. EC
Immature EC
VWF EC
AEC
Prolif. AEC
HE
YS
LR probability predicted
1.0
0.0
Prolif. Sin. EC
Immature EC
VWF EC
AEC
HE
EC
Immature EC
Sinusoidal EC
Prolif. EC
tip EC
Sinusoidal EC
Prolif. Sin. EC
Immature EC
VWF EC
AEC
HE
Sinusoidal EC
Prolif. Sin. EC
Immature EC
AEC
HE
AGM
EL
FBM
Definitive iPSC
iPSC
C
iPSC (Definitive)
AEC
HE
HSPC 1
HSPC 2
FDG2
FDG1
n=2; k=900
IL33
ALDH1A1
log count
YS
iPSC
iPSC (Definitive)
KCNK17
SPINK2
n=3; k=2,262
n=6; k=368
D
EFNB2/EPHB4
TNFSF10/RIPK1
PDGFB/PDGFRB
JAG1/NOTCH2
TGFB2/TBGbeta receptor 1
IL35/IL35 receptor
DLL1/NOTCH2
CSF3/CSF3R
PLD2/ARF1
EPO/EPOR
EPO/EFNB2
BMP7/PTPRK
BMP8B/PLAUR
WNT4/SMO
BMP10/VSIR
WNT7B/FZD4
GDF9/TGFR/BMPR2
TGFB3/TGFBR3
PLA2G2A/a5b1 complex
TGFB3/TGFbeta receptor 1
PLA2G2A/a4b1 complex
IL13/TMEM219
CD94:NKG2E/HLA-E
LEP/LEPR
CD70/GPRC5B
EC↓
Endoderm↓
Fibroblast↓
SM↓
Endoderm↑
Fibroblast↑
Fibroblast/SM↑
~4 PCW
~5 PCW
~7 PCW
~8 PCW
mean z-score
-1 0 1
AEC/HSPC
Sin. EC/HSPC
VWF EC/HSPC
Endoderm/HSPC
Fibroblast/HSPC
SM/HSPC
E
AEC/HSPC
Prolif. Sin. EC/HSPC
Endoderm/HSPC
Fibroblast/HSPC
Smooth Muscle/HSPC
AGM
Endothelium/HSC
Hepatocyte/HSC
Fibroblast/HSC
EL
mean z-score
-1 0 1
F
Adhesion
AEC/HSPC
Endoderm/HSPC
Fibroblast/HSPC
Prolif. Sin. EC/HSPC
Smooth Muscle/HSPC
AGM
Endothelium/HSC
Hepatocyte/HSC
Fibroblast/HSC
EL
AEC/HSPC
Sin. EC/HSPC
VWF EC/HSPC
Endoderm/HSPC
Fibroblast/HSPC
Smooth Muscle/HSPC
YS
~4 PCW
~5 PCW
~7 PCW
~8 PCW
mean z-score
-1 0 1
SELP/CD34
SELP/SELPLG
SELL/CD34
SELL/SELPLG
JAM2/a4b1 complex
VCAM1/a4b1 complex
VCAM1/a9b1 complex
COL18A1/a1b1 complex
COL4A1/a1b1 complex
COL9A2/a1b1 complex
COL8A2/a1b1 complex
COL8A1/a1b1 complex
COL2A1/a1b1 complex
G
Primitive
HSPC
HSPC (C.)
~4 PCW
~5 PCW
~7 PCW
~8 PCW
Mean z-score expression
1.5
1.0
0.5
0.0
0.5
1.0
1.5
Oxidative Resp.
Cell Cycle
Apoptosis
Definitive
HSPC
HSPC (C.)
~5 PCW
~7 PCW
~8 PCW
Mean z-score expression
1.0
0.5
0.0
0.5
1.0
1.5
Oxidative Resp.
Cell Cycle
Apoptosis

**Fig. S7.**

**Fig. S7: The lifespan of YS HSPCs**

**(A)** Dot plot showing the expression level (colour scale) and percent expression (dot size) of genes distinguishing endothelial cell subsets in the YS (main) scRNA-seq data, as well as the matched EL scRNA-seq and the iPSC scRNA-seq dataset (*21*).

(**B**) Median logistic regression class projection probabilities for a model trained on YS endothelial cell (EC) states (y-axis) projected onto EC states in AGM (*12*, *58*), matched EL, and FBM (*54*), iPSC (*21*) and definitive iPSC (*11*) (x-axis) (vmin=0, vmax=1) (**Table S16**).

**(C)** Top: FDG overlaid with PAGA showing trajectory of HE transition to HSPC in definitive iPSC scRNA-seq dataset (*11*) (n=3, k=2,262) with feature plots of key genes (*IL33*, *ALDH1A1*) involved in endothelial to hemogenic transition (**Table S5**). Bottom: Feature plots of key genes in endothelial to hemogenic transition (S*PINK2, KCNK17*) in YS, iPSC and definitive iPSCs scRNA-seq trajectories (also shown in **Fig. 4C**).

**(D)** Dot plot showing the level (colour scale) and percent expression (dot size) of genes predicted by CellphoneDB to form statistically significant interactions between YS HSPC/ cycling HSPC and ECs, fibroblasts, smooth muscle cells, and endoderm. Brackets indicate genes which form complexes (data scaled max_value=10) (**Table S24**).

**(E)** Heatmap showing relative mean expression z-scores of curated and statistically significant ($p<0.05$) CellphoneDB putative receptor ligand interactions between AGM (top) and EL (bottom) stromal subsets vs HSPC across gestation. Growth factor, TGF beta and NOTCH ligand receptor -related gene interactions have been highlighted (**Table S24**).

**(F)** Heatmap showing relative mean expression z-scores of curated and statistically significant ($p<0.05$) CellphoneDB putative curated functional adhesion receptor ligand interactions between AGM (top), YS (middle) and EL (bottom) stromal subsets vs HSPC across gestation (**Table S24**).

**(G)** Heatmaps showing mean z-scored expression of metabolic (GO-ontology, GO-0045333), cell cycle (GO-ontology, GO:0022402) and apoptosis (GO-ontology GO:0006915) modules for primitive (left) and definitive (right) YS HSPC and cycling HSPC across gestational age.

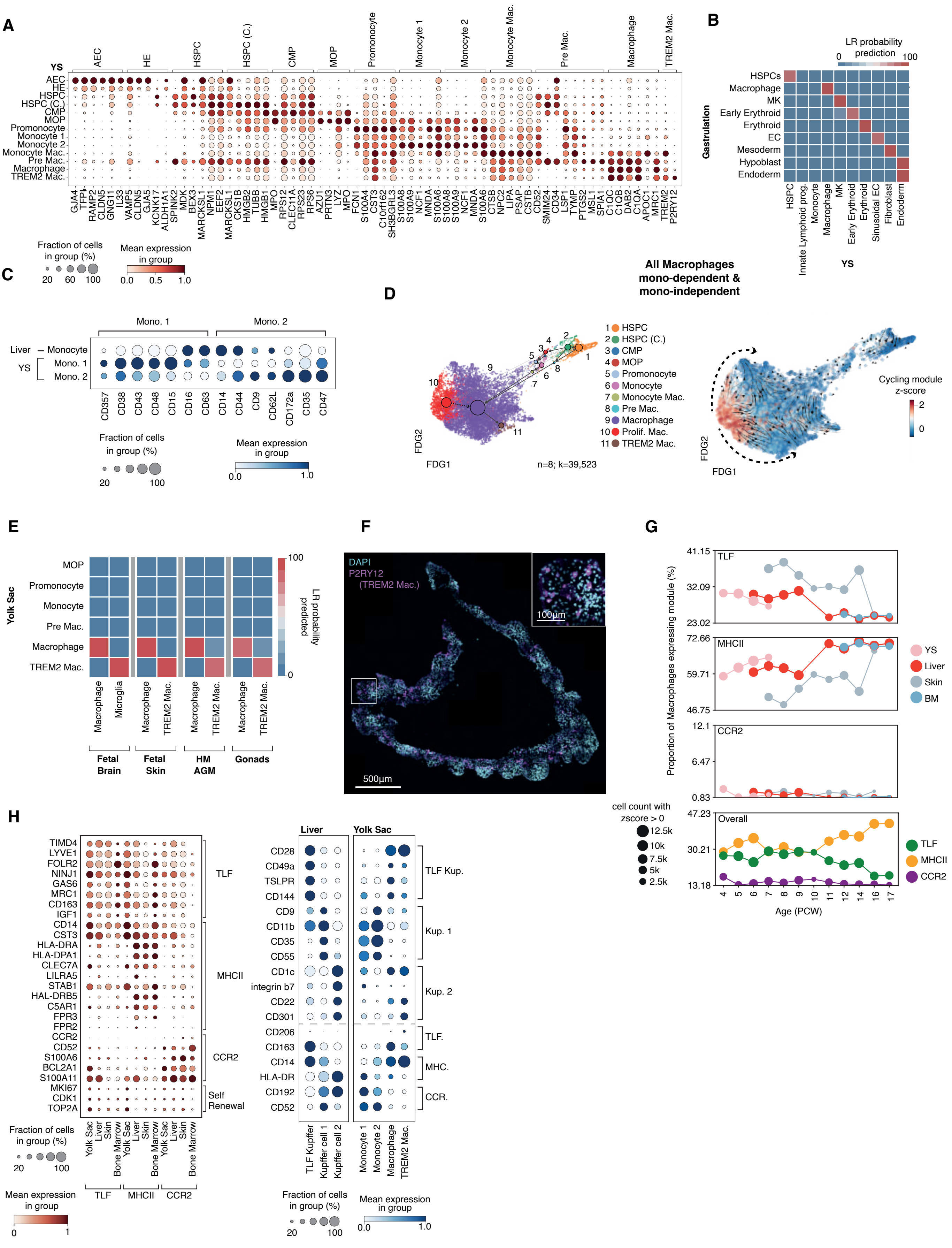

A
YS
AEC
HE
HSPC
HSPC (C.)
CMP
MOP
Promonocyte
Monocyte 1
Monocyte 2
Monocyte Mac.
Pre Mac.
Macrophage
TREM2 Mac.
GJA4 TFPI RAMP2 CLDN5 GNG11 IL33 VAMP5 CLDN5 GJA5 KCNK17 ALDH1A1 SPINK2 MDK BEX3 MARCKSL1 NPM1 EEF2 MARCKSL1 CKS1B HMGB2 TUBB HMGB1 MPO RPS12 CLEC11A RPS23 RPS6 AZU1 PRTN3 LYZ MPO FCN1 S100A4 CST3 C10rf162 SH3BGRL3 S100A8 S100A9 NCF1 MNDA S100A6 S100A8 S100A9 NCF1 MNDA S100A6 CTSB NPC2 LIPA PSAP CSTB CD52 SMIM24 CD34 LSP1 TYMP PTGS2 MSL1 SPIA1 C1QC C1QB DAB2 C1QA APOC1 MRC1 TREM2 P2RY12
Fraction of cells in group (%)
20 60 100
Mean expression in group
0.0 0.5 1.0
B
LR probability prediction
0 100
Gastrulation
HSPCs
Macrophage
MK
Early Erythroid
Erythroid
EC
Mesoderm
Hypoblast
Endoderm
HSPC
Innate Lymphoid prog.
Monocyte
Macrophage
MK
Early Erythroid
Erythroid
Sinusoidal EC
Fibroblast
Endoderm
YS
C
Mono. 1
Mono. 2
Liver – Monocyte
YS
Mono. 1
Mono. 2
CD357 CD38 CD43 CD48 CD15 CD16 CD63 CD14 CD44 CD9 CD62L CD172a CD35 CD47
Fraction of cells in group (%)
20 100
Mean expression in group
0.0 1.0
All Macrophages mono-dependent & mono-independent
D
1 HSPC
2 HSPC (C.)
3 CMP
4 MOP
5 Promonocyte
6 Monocyte
7 Monocyte Mac.
8 Pre Mac.
9 Macrophage
10 Prolif. Mac.
11 TREM2 Mac.
FDG2
FDG1
n=8; k=39,523
Cycling module z-score
2
0
E
Yolk Sac
MOP
Promonocyte
Monocyte
Pre Mac.
Macrophage
TREM2 Mac.
Macrophage
Microglia
Macrophage
TREM2 Mac.
Macrophage
TREM2 Mac.
Macrophage
TREM2 Mac.
Fetal Brain
Fetal Skin
HM AGM
Gonads
100
0
LR probability predicted
F
DAPI
P2RY12 (TREM2 Mac.)
100µm
500µm
G
Proportion of Macrophages expressing module (%)
TLF
41.15
32.09
23.02
MHCII
72.66
59.71
46.75
CCR2
12.1
6.47
0.83
Overall
47.23
30.21
13.18
YS
Liver
Skin
BM
cell count with zscore > 0
12.5k
10k
7.5k
5k
2.5k
TLF
MHCII
CCR2
4 5 6 7 8 9 10 11 12 14 16 17
Age (PCW)
H
TIMD4 LYVE1 FOLR2 NINJ1 GAS6 MRC1 CD163 IGF1
TLF
CD14 CST3 HLA-DRA HLA-DPA1 CLEC7A LILRA5 STAB1 HAL-DRB5 C5AR1 FPR3 FPR2
MHCII
CCR2 CD52 S100A6 BCL2A1 S100A11
CCR2
MKI67 CDK1 TOP2A
Self Renewal
Yolk Sac Liver Skin Bone Marrow Yolk Sac Liver Skin Bone Marrow Yolk Sac Liver Skin Bone Marrow
TLF
MHCII
CCR2
Fraction of cells in group (%)
20 100
Mean expression in group
0 1
Liver
Yolk Sac
CD28 CD49a TSLPR CD144
TLF Kup.
CD9 CD11b CD35 CD55
Kup. 1
CD1c integrin b7 CD22 CD301
Kup. 2
CD206
TLF.
CD163 CD14
MHC.
HLA-DR CD192 CD52
CCR.
TLF Kupffer
Kupffer cell 1
Kupffer cell 2
Monocyte 1
Monocyte 2
Macrophage
TREM2 Mac.
Fraction of cells in group (%)
20 100
Mean expression in group
0.0 1.0

**Fig. S8.**

**Fig. S8: Accelerated macrophage production in YS and iPSC**

**(A)** Dot plot showing the level (colour scale) and percent expression (dot size) of myeloid lineage gene markers in YS scRNA-seq dataset (**Table S3**).

**(B)** Median logistic regression class prediction probabilities for a model trained on YS scRNA-seq cell states (x-axis) and projected onto cell states in human gastrulation scRNA-seq data (*57*) (y-axis). This LR was performed after reannotating human gastrula data in-house (**Table S17**).

**(C)** Dot plot showing the level (colour scale) and percent expression (dot size) of proteins differentially expressed and matched to RNA markers shown in **Fig. 5A** per matched embryonic liver Monocytes, YS Monocyte1 and YS Monocyte2 in respective CITE-seq datasets (**Table S25**).

**(D)** Left: FDG overlaid with directional PAGA showing both trajectory progenitor-derived macrophages and proliferating macrophages contributing to macrophage differentiation from CS10-CS23 (n=8, k=39,523). Right: CellRank state transition matrix inferred arrows projected onto FDG indicate the trend of trajectory, and colour shows z-score enrichment in cycling module (GO:0007049) genes showing contribution of proliferating macrophages to macrophage population.

**(E)** Median logistic regression class prediction probabilities for a model trained on YS scRNA-seq myeloid cell states projected onto fetal brain (*44*), fetal skin (*45*), AGM (*11*) and testes (*46*) external scRNA-seq datasets with a classification probability threshold of >0.7 showing distinct cross-tissue matched macrophage and microglia/TREM2 macrophage cell state profiles (vmax=1, vmin=-1) (**Table S18**).

**(F)** Immunofluorescence microscopy staining of P2RY12 in n=1 YS at 8PCW. Scale bar=500um (100um, inset).

**(G)** Line graph showing the relative change proportion of YS scRNA-seq macrophages (y-axis) enriched in expression of TLF, MHCII and CCR2 expression modules (z-scored module score > 0) over YS (pink), liver (red), skin (grey) and BM (blue) across gestational age, including *TIMD4, LYVE1,FOLR2* for TLF macrophages, CD14, CST3, HLA-DRA for MHCII high macrophages and CCR2, CD52, S100A6 for CCR2 high macrophages. Size of spots indicates the number of cells expressing each module with a z-score>0.

**(H)** Left: Dot plot showing the scaled (colour scale)(max value =1) and percent expression (dot size) of TLF, MHCII and CCR2 expression modules in YS, Liver, Skin and Bone-marrow scRNA-seq macrophages (x-axis). Right: Dotplot of differentially expressed Cite-seq proteins across TLF Kupffer cells, Kupffer1 and Kupffer 2 in embryonic liver data, Monocyte 1, Monocyte 2, Macrophages and Trem2 Macrophages in YS data (**Table S27**).

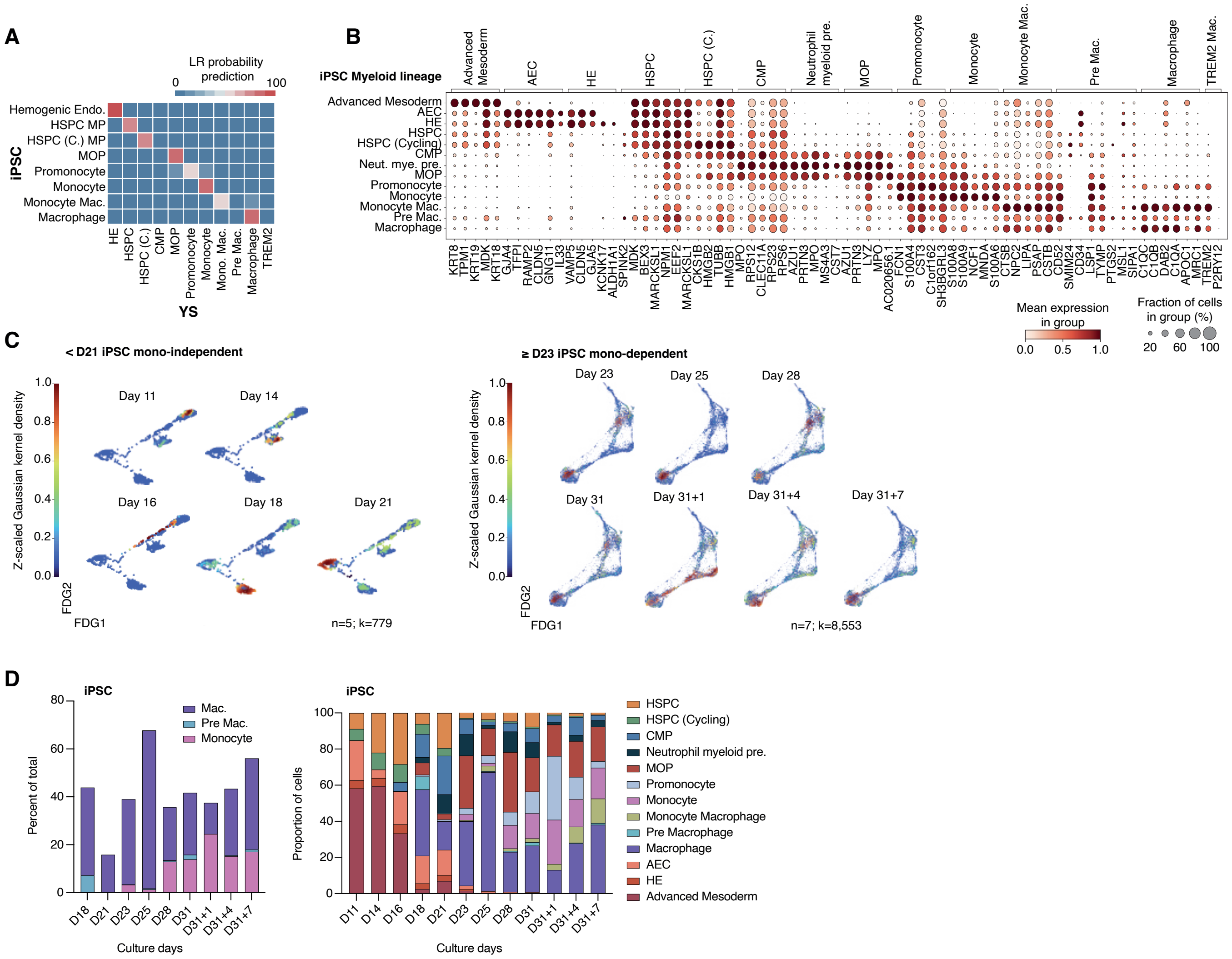
A
B
C
D
LR probability prediction
iPSC
YS
iPSC Myeloid lineage
Mean expression in group
Fraction of cells in group (%)
< D21 iPSC mono-independent
≥ D23 iPSC mono-dependent
Z-scaled Gaussian kernel density
n=5; k=779
n=7; k=8,553
Percent of total
Proportion of cells
Culture days
HSPC
HSPC (Cycling)
CMP
Neutrophil myeloid pre.
MOP
Promonocyte
Monocyte
Monocyte Macrophage
Pre Macrophage
Macrophage
AEC
HE
Advanced Mesoderm

**Fig. S9.**

**Fig. S9: Accelerated macrophage production in YS and iPSC**

**(A)** Median logistic regression class prediction probabilities for a model trained on YS scRNA-seq myeloid cell states (x-axis) projected onto equivalent cell states in iPSC scRNA-seq data (y-axis) (*21*).'Monocyte1' and 'Monocyte2' are grouped together into the 'Monocyte' category **(Table S14)**.

**(B)** Dot plot showing the level (colour scale) and percent expression (dot size) of myeloid lineage gene markers in iPSC scRNA-seq (*21*) dataset myeloid-lineage cell states (**Table S7**).

**(C)** Density plots showing the distribution of transitioning iPSC-derived scRNA-seq (*21*) macrophage lineage cells from <D21 (n= 5; k=779) (left) >D23 (n=7; k=8,553) (right) in the integrated FDG landscape. Colour of cells represents the z-scored kernel density estimation (KDE) score for each timepoint (**Table S7, S5)**.

**(D)** Left: Stacked bar plot displaying the percent of monocytes, pre-macrophages and macrophages found in iPSC cultures (*21*) by day. Colours match cell states in **c**. Right: Stacked bar plot displaying the proportion of cell states found in iPSC cultures by day.

**Movie S1. (separate file)**

Light-sheet fluorescence microscopy video of ~6.9 PCW YS stained with CD34 (yellow), HNF4A (cyan), and LYVE1 (magenta). Stillshot taken for the left panel of Figure 1D.

**Movie S2. (separate file)**

Light-sheet fluorescence microscopy video of ~5.7 PCW YS stained with ECAD (yellow), HNF4A (cyan), and LYVE1 (magenta). Stillshot taken for the right panel of Figure 1D.

**Movie S3. (separate file)**

Light-sheet fluorescence microscopy video of ~7 PCW YS stained with PLVAP (green), LYVE1 (red), and IBA1 (white). Stills taken for the Supplementary Figure 3F.

**Table S1. (seperate file)**

Sample metadata: The sample information metadata for the core yolk sac and embryonic liver single cell datasets (scRNA-seq, CITE-seq and Smart-seq2) that have been integrated and analysed for this study. Sample metadata includes, among other categories, sample ID, age, sex, tissue, sequencing platform, and alignment sofware. Some data has previously been published by ourselves together with our collaborators - this is noted clearly in the dataset column. All data included are made publicly available on ArrayExpress at the accessions noted, inclusive of raw FASTQ files and raw count matrices.

**Table S2. (seperate file)**

Sample manifest: The sample manifest which directly follows the sample metadata table. This manifest includes summaries for the yolk sac and embryonic liver scRNA-seq, CITE-seq and Smart-seq2 data that has been integrated and analysed for this study. Metadata and cell counts information are provided more broadly for each biological replicate and tissue.

**Table S3. (seperate file)**

Yolk sac scRNA-seq (10x) metadata: Metadata for the yolk sac 10x scRNA-seq data including scrublet QC calls by barcode.

**Table S4. (seperate file)**

Yolk sac and liver scRNA-seq and CITE-seq (10x) cell state counts:
Cell state annotation counts in each of the core yolk sac and embryonic liver single cell datasets (scRNA-seq, CITE-seq and Smart-seq2) desegregated by biological replicate.

**Table S5. (seperate file)**

Coordinates metadata: Dimensional reduction coordinates by cell barcode for all UMAP and FDG embeddings shown in the main and supplementary figures of this study.

**Table S6. (seperate file)**

External datasets: A table detailing all external single cell datasets leveraged for comparison with our core human yolk sac and embryonic liver single cell datasets. For each external dataset, we note organ, research article doi, data accessibility portal links, number of biological replicates and cells, and provide any notes required for subsetting.

**Table S7. (seperate file)**

External Dataset annotations: Barcodes cell annotations for all external datasets used which are present within both main and extended figure panels shown.

**Table S8. (seperate file)**

Smart-Seq2 Metadata: Metadata for the yolk sac Smart-seq2 data by barcode.

**Table S9. (seperate file)**

Yolk sac CITE-seq metadata: Metadata for the yolk sac CITE-seq data by barcode for RNA and protein.

**Table S10. (seperate file)**
Logistic Regression EL scRNA-seq to Liver Citeseq: Table output from the Logistic regression ElasticNet model trained on the combined low-dimensional representation of the EL scRNA-seq data for purposes of projection, class assignment and visualisation of class correspondence between trained labels and target YS scRNA-seq data annotations. (See methods Cell state predictions using probabilistic low-dimensional ElasticNet regression for default parameters used)

**Table S11. (seperate file)**
Logistic Regression YS scRNA-seq to YS ss2 scRNA-seq: Table output from the Logistic regression ElasticNet model trained on the combined low-dimensional representation of the YS scRNA-seq data for purposes of projection, class assignment and visualisation of class correspondence between trained labels and target YS SS2 scRNA-seq plate-based data annotations. (See methods Cell state predictions using probabilistic low-dimensional ElasticNet regression for default parameters used)

**Table S12. (seperate file)**
Logistic Regression YS scRNA-seq to EL scRNA-seq: Table output from the Logistic regression ElasticNet model trained on the combined low-dimensional representation of the YS scRNA-seq data for purposes of projection, class assignment and visualisation of class correspondence between trained labels and target EL RNA data annotations. (See methods for default parameters used)

**Table S13. (seperate file)**
Logistic Regression YS scRNA-seq to mouse yolk sac scRNA-seq: Table output from Logistic regression showing the mean probability scoring trained using YS scRNA-seq data used to classify individual cell states within mouse yolk sac scRNA-seq. Clusters were then assigned classes by the majority projected label that had a label count of > (mean + (1*std)) of label counts per cluster.

**Table S14. (seperate file)**
Logistic Regression YS scRNA-seq to iPSC scRNA-seq: Table output from the Logistic regression ElasticNet model trained on the combined low-dimensional representation of the YS scRNA-seq data for purposes of projection, class assignment and visualisation of class correspondence between trained labels and target iPSC scRNA-seq data annotations. (See methods Cell state predictions using probabilistic low-dimensional ElasticNet regression for default parameters used)

**Table S15. (seperate file)**
Logistic Regression YS scRNA-seq to mouse hematopoietic lineage: Table output from the Logistic regression ElasticNet model trained on the combined low-dimensional representation of the cross species SAMAP sequence-blast YS scRNA-seq data for purposes of projection, class assignment and visualisation of class correspondence between trained labels and target mouse hematopoietic lineage scRNA-seq data annotations. (See methods Cell state predictions using probabilistic low-dimensional ElasticNet regression for default parameters used)

**Table S16. (seperate file)**

Logistic Regression YS scRNA-seq to AGM, EL and FBM endothelium: Table output from the Logistic regression ElasticNet model trained on the combined low-dimensional representation of the YS scRNA-seq data for purposes of projection, class assignment and visualisation of class correspondence between trained labels and target datasets AGM, EL and FBM RNA-seq data annotations. (See methods for default parameters used)

**Table S17. (seperate file)**

Logistic regression projection probabilities for Yolksac Gastrulation SS2 data: Table output from the Logistic regression ElasticNet model trained on the combined low-dimensional representation of the YS scRNA-seq data for purposes of projection, class assignment and visualisation of class correspondence between trained labels and target YS gastrulation SS2 data annotations. (See methods for default parameters used)

**Table S18. (seperate file)**

Logistic regression projection probabilities for EC populations across organs: Table output from the Logistic regression ElasticNet model trained on the combined low-dimensional representation of the YS scRNA-seq data for purposes of projection, class assignment and visualisation of class correspondence between trained labels and target fetal brain and fetal skin myeloid lineage scRNA-seq data annotations. (See methods Cell state predictions using probabilistic low-dimensional ElasticNet regression for default parameters used)

**Table S19. (seperate file)**

MILO differential abundance analysis output: A table containing differentially abundance of cell state neighbourhoods by MILO with differential expression testing results across each cell state enriched in early and late neighbourhoods. Significance of differential abundance was taken as SpatialFDR (<0.1, logFC <0) for early enriched neighbourhoods and SpatialFDR (<0.1, logFC >0) for later enriched neighbourhoods. (See methods for Differential abundance testing and FACs correction).

**Table S20. (seperate file)**

EL scRNA-seq (10X) metadata: Metadata for the embryonic liver 10x scRNA-seq data by barcode.

**Table S21. (seperate file)**

Conserved and differential clustered enriched gene set modules statistics between Mouse and Human endoderm scRNA-seq (10X): Neighbourhood clusters and significance information of significantly enriched gene set modules for differential and conserved expression between cell states of interest (See methods for Clustered gene-set enrichment analysis).

**Table S22. (seperate file)**

GSEA enrichment statistics: Gene set enrichment outputs produced by the enrichr (P values provided from Fisher exact test and ranked by z-score to expected background enrichment) workflow as implemented in the GSEApy package (See methods for gene set enrichment analysis).

**Table S23. (seperate file)**
CellphoneDB receptor-ligand interactions statistics across YS scRNA-seq cellstates data (10X): Significant CellPhoneDB receptor ligand interaction predictions between all cellstates in the YS scRNA-seq data ran using the CellPhoneDB statistical method workflow (see methods for Cell-cell interaction predictions using CellPhoneDB). Significance threshold for mean receptor ligand expression was taken as ($P<0.05$).

**Table S24. (seperate file)**
CellphoneDB receptor-ligand interactions statistics between HSPCs and Stroma in YS, EL and AGM scRNA-seq cellstates data (10X): Significant CellPhoneDB receptor ligand interaction predictions between HSPC cellstates and stroma in the YS,EL and AGM scRNA-seq data desegregated by gestational time ran using the CellPhoneDB statistical method workflow (see methods for Cell-cell interaction predictions using CellPhoneDB). Significance threshold for mean receptor ligand expression was taken as ($P<0.05$).

**Table S25. (seperate file)**
Differentially expressed proteins in CITE-seq datasets (YS/EL) monocyte fractions: Differentially expression testing results from independent analyses between monocyte subsets in the YS CITE-seq dataset derived from the sc.tl.rank_gene_groups function in the Scanpy package which performed a two-sided Wilcoxon rank-sum test. P-values significance threshold was taken as $P <0.05$.

**Table S26. (seperate file)**
pyScenic differential regulon usage stats: Table containing differential regulon usages across the YS scRNA-seq Macrophage differentiation pseudotime. Significantly changing regulons across pseudotime were estimated by a General additive model (See methods for psScenic) (P_value < 0.05). Significant GAMs with a logFC +/- 0 for each cell state were retained for ranking in visualisation.

**Table S27. (seperate file)**
Differentially expressed proteins in Liver CITE-seq Kupffer cells: Differentially expression testing results from independent analyses between Kupffer subsets in the Liver CITE-seq dataset derived from the sc.tl.rank_gene_groups function in the Scanpy package which performed a two-sided Wilcoxon rank-sum test. P-values significance threshold was taken as $P <0.05$.

**Table S28. (seperate file)**
Differentially expressed genes for scRNA-seq datasets: Differentially expression testing results from independent analyses between cell states in the YS, EL and IPSC scRNA-seq datasets derived from the sc.tl.rank_gene_groups function in the Scanpy package which performed a two-sided Wilcoxon rank-sum test for genes expressed in >25% of cells, log-transformed fold change cut-off of 0.25. All p-values were adjusted for multiple testing using the Benjamini–Hochberg method and significance threshold was taken as $P <0.05$.

**Table S29. (seperate file)**
CITE-seq antibody details: Details of the TotalSeq™ (Biolegend) antibodies, including target, clone and barcode, in the CITE-seq cocktail used on embryonic YS and liver.

**Table S30. (seperate file)**

Liver CITE-seq metadata: Metadata for the Liver CITE-seq data by barcode for RNA and protein.

**Table S31. (seperate file)**

Logistic Regression YS scRNA-seq to YS CITE-seq: Table output from the Logistic regression l2 model (sparcity=0.2, max_iter=1000) trained on the combined low-dimensional representation of the YS scRNA-seq data for purposes of projection, class assignment and visualisation of class correspondence between trained labels and target YS Cite-seq RNA data annotations. (See methods for Cell state predictions using probabilistic low-dimensional ElasticNet regression)

**Table S32. (seperate file)**

Antibodies Table: List of antibodies used for cell sorting and imaging: i) Enrichment sort for scRNA-seq (10x, CITE-seq and SS2), ii) IHC, iii) FFPE immunofluorescence microscopy, iv) Fixed frozen immunofluorescence microscopy, v) Light-sheet fluorescence microscopy, vi) RNAscope-associated immunofluorescence microscopy
